# A metabolic network-based approach for developing feeding strategies for CHO cells to increase monoclonal antibody production

**DOI:** 10.1101/751347

**Authors:** Hamideh Fouladiha, Sayed-Amir Marashi, Fatemeh Torkashvand, Fereidoun Mahboudi, Nathan E. Lewis, Behrouz Vaziri

**Author notes:** Corresponding authors: Dr. Sayed-Amir Marashi, E-mail address, Dr. Behrouz Vaziri.

## Abstract

Chinese hamster ovary (CHO) cells are the main workhorse in the biopharmaceutical industry for the production of recombinant proteins, such as monoclonal antibodies. To date, a variety of metabolic engineering approaches have been used to improve the productivity of CHO cells. While genetic manipulations are potentially laborious in mammalian cells, rational design of CHO cell culture medium or efficient fed-batch strategies are more popular approaches for bioprocess optimization. In this study, a genome-scale metabolic network model of CHO cells was used to design feeding strategies for CHO cells to improve monoclonal antibody (mAb) production. A number of metabolites, including threonine and arachidonate, were suggested by the model to be added into cell culture medium. The designed composition has been experimentally validated, and then optimized, using design of experiment methods. About a two-fold increase in the total mAb expression has been observed using this strategy. Our approach can be used in similar bioprocess optimization problems, in order to suggest new ways of increasing production in different cell factories.

## Introduction

Bio-based production of industrially-relevant or pharmaceutically-important proteins, such as monoclonal antibodies (mAbs), has been a major goal of biotechnology in the past decades [1]. MAbs have diverse diagnostic and therapeutic applications, especially for cancer and autoimmune diseases [2]. The existence of certain glycosylation patterns on some mAbs restricts the choice of host cell lines for their production to mammalian cell lines, including Chinese hamster ovary (CHO) cells.

Several studies have focused on optimizing the production of recombinant proteins in CHO cells [3-5]. In such studies, different omics approaches were used for understanding the reasons of higher levels of mAb production in a selected CHO cell line [6-8]. Several genetic manipulation methods have been used to design super-producer CHO cells [9]. However, such methods can be potentially laborious for mammalian cells. Therefore, bioprocess optimization methods, including rational design of feeding strategies or finding optimized composition for cell culture medium, are attractive for increasing productivity of CHO cells [10-12].

Growth and productivity of CHO cells are under the direct influence of cell metabolism [13]. To simulate the metabolism of cells, genome-scale metabolic network models (GEMs) can be very useful. GEMs represent *in silico* models of metabolism. A standard GEM includes organism-specific biochemical reactions together with accurate gene-protein-reaction (GPR) relations. Using reliable metabolic models, one can perform virtual experiments in a rapid and cheap manner [14]. Therefore, GEMs are considered as helpful tools in cell biology and metabolic engineering, because of their potential for predicting the metabolic state of cells under certain growth conditions [15].

In this study, we proposed a new approach for fermentation process optimization, in which a metabolic modeling study was used upstream of the design of experiment (DoE) methods as a means to reduce the number of variables in cell culture medium optimization. Here, a GEM of CHO cells, called *i*CHO1766 [16], has been used in order to predict strategies for increasing mAb production in CHO cells. Several constraint-based modeling methods have been developed to design new cell factories [17]. In our study, the FVSEOF method [18] was used to suggest experimental ways of increasing the production of a mAb in CHO cells (<u>F</u>lux <u>V</u>ariability <u>S</u>canning based on <u>E</u>nforced <u>O</u>bjective <u>F</u>lux or <u>FVSEOF</u>). Using FVSEOF method, it was predicted that CHO cell culture medium supplementation with 15 metabolites might be useful for increasing mAb production. Then, *in vitro* experiments were designed by DoE methods to screen these 15 metabolites, to find the most effective metabolites for increasing mAb production. To the best of our knowledge, this is the first time that a GEM of CHO cells has been used to design a feeding strategy. In addition, among the predicted 15 metabolites used in the feeds, there are some metabolites that have not been reported as supplements in CHO cell culture ever before.

## Materials and methods

### Cell culture

The CHO cell line, producing an anti-α4β7 antibody, was gifted from Radin Biotech Company, Iran. The cells were cultivated in proCHO5 cell culture medium (purchased from Lonza AG, Verviers, Belgium), supplemented with 4 mM L-glutamine, 0.1% of anti-clumping agent, 1% of pluronic F68 and 1% of Pen-Strep (all purchased from Gibco, Life Technology, USA). Anti-clumping agent is a chemically formulated solution to inhibit the cells from sticking to each other in the cell culture medium. It is an animal-free solution and does not contain any enzymes or proteins.

CHO cells were cultured in 20 ml glass bottles (Duran Schott®) with a working volume of 5 ml. The bottles were placed in a shaker that was installed in a CO2 incubator to be agitated mechanically at 120 rpm. The cells were incubated at 37°C with a 5% solution of CO_2_. Each vessel was inoculated with a cell density of 5×10^5^ viable cells/ml, when the viability was more than 80%. The cells were cultivated in fed-batch mode and were supplemented with 0.5 mL of the feed solutions on day 3, 5, 7, 9, and 11.

Feed solutions include different compositions of 15 metabolites dissolved in proCHO5 cell culture medium. Amino acid powders were purchased from HiMedia Laboratories (Mumbai, India). Vitamins and other metabolites were purchased from Sigma-Aldrich Company (Germany). All metabolites were dissolved in proCHO5 cell culture medium.

Cell viability was determined daily by Trypan Blue assay, using a Neubauer cytometer. Every day, the cell culture bottles were carefully transferred to a laminar hood. Then, 100 microliters of the working volume were transferred to a microtube using a sampler. The bottles were back in the CO_2_ incubator. The integral viable cell count (IVCC) was calculated by cumulative addition of viable cell count in each day of cell culture. The cells were harvested when the viabilities dropped to less than 50%. In the harvesting day, the cell culture medium was centrifuged in 1100 rpm for 5 minutes, and the supernatant was 0.2 µm filtered and stored in −20 °C for determination of mAb concentration.

### Metabolic network modeling

A genome-scale metabolic network model of CHO cells, called *i*CHO1766 [16], was used in our study. Different formats of *i*CHO1766 are included in the supplementary file of its article in the following link (http://dx.doi.org/10.1016/j.cels.2016.10.020). In addition, *i*CHO1766 can be browsed and downloaded at http://www.CHOgenome.org and at the BiGG Models database (http://bigg.ucsd.edu). The metabolic reaction of mAb production was added to the model, according to the amino acid composition of the mAb which is produced by CHO cells *in vitro*.

The reaction for constructing the heavy chain of the mAb:

34 Pro + 901 H2O + 452 ATP + 4 Met + 33 Gly + 13 Phe + 22 Glu + 20 Ala + 20 Asn + 11 Cys + 16 Gln + 55 Ser + 34 Thr + 12 Arg + 16 Asp + 12 His + 33 Leu + 10 Ile + 31 Lys + 45 Val + 10 Trp + 20 Tyr + 900 GTP => 901 H^+^ + ADP + 901 Pi + 900 GDP + 451 AMP + 451 PPi + Heavy_Chain.

The reaction for constructing the light chain of the mAb:

11 Pro + 435 H2O + 219 ATP + Met + 13 Gly + 8 Phe + 11 Glu + 13 Ala + 6 Asn + 5 Cys + 15 Gln + 33 Ser + 17 Thr + 6 Arg + 13 Asp + 3 His + 15 Leu + 6 Ile + 13 Lys + 15 Val + 2 Trp + 12 Tyr + 434 GTP => 435 H^+^ + ADP + 435 Pi + 434 GDP + 218 AMP + 218 PPi + Light_Chain.

The reaction for final formation of the mAb and addition of the glycans:

2 N_linked_glycans + 2 Heavy_Chain + 2 Light_Chain => mAb

FVSEOF algorithm [18] was used to predict strategies for increasing the production of a mAb in CHO cells. In brief, using the FVSEOF method and the metabolic network, the effects of an enforced theoretical increase in mAb production on the rates of all other metabolic reactions of CHO cells can be modeled. Then, a list of metabolic reactions is generated as FVSEOF output (Supporting information, S0 file). This list includes the metabolic reactions that have altered rates during the theoretical increase in mAb production. FVSEOF suggest that *in vitro* changes in the reactions of this list, according to the direction of their predicted altered rates in the list, can increase mAb production *in vitro*. The in-house implementation of FVSEOF algorithm was done using COBRA Toolbox [19].

### Design of Experiment (DoE) methods

The Plackett-Burman (PB) design, Central Composite Design (CCD), and statistical analysis of the results were performed using the Design-Expert software version 7 (Stat-EaseInc. Minneapolis, Minnesota, USA). Two responses were defined for both the PB design and CCD. The first response was the integral viable cell count (IVCC) of the harvesting day. IVCC of each day was calculated by cumulative addition of viable cell counts (from the first day of cell culture until the end of each day). The second response was the total mAb expression of each group. This response was calculated by measuring the concentration of mAb in the CHO cell culture supernatant on the harvesting day, measured by HPLC (see the next section of methods).

### MAb concentration determination

A MAbPac protein A affinity column (Thermo-Scientific, USA) was used for HPLC analysis. The equilibration and elution buffers have the same compositions, which includes 5% of PBS buffer and 5% acetonitrile in 0.15 M sodium chloride solution (Merck, Darmstadt, Germany). The pH of equilibration buffer was 7.2, and the pH of elution buffer was set to 2.7 using ortho-phosphoric acid. The temperature for the column was set to 30°C. The relative concentration of elution buffer to equilibration buffer was increased from 0 to 100% in approximately 3 minutes. The injection volume was 20 µL, and the flow rate was 2 ml/minute. The concentration of mAb in CHO cell culture medium was determined based on a standard curve generated with previously known concentrations of a purified monoclonal IgG as the standard solution.

## Results

In our study, a metabolic model of CHO cells, named *i*CHO1766, and a computational modeling method, named FVSEOF, were used to design feeding strategies to increase mAb production. The number of reactions and genes of *i*CHO1766 is relatively high, that limits the number of computational methods of strain design available to be tested on the model. Therefore, in our study, the FVSEOF method was used. The previous version of the FVSEOF method had been successfully used to suggest experimental ways of increasing a recombinant protein production in *Pichia pastoris* [20]. The output of the FVSEOF method is a list of reactions that are suggested to be changed *in vitro* to increase mAb production. In our study, the exchange reactions of this list that had an increased consumption rate were selected to be tested *in vitro*. These selected reactions were related to the exchange of 15 metabolites, including 7 amino acids (glutamine, asparagine, lysine, tryptophan, threonine, valine, and histidine), 3 vitamins (vitamin A, B1 and B6), and 5 other metabolites (thymidine, deoxy-cytidine, 3-methyl-oxobutyrate, deoxy-guanosine, and arachidonate). It should be noted that we restricted our analysis only to exchange reactions. Therefore, these reactions presumably belong to separate metabolic pathways or groups. In other words, these reactions are not coupled to each other. In addition to exchange reactions, some internal metabolic reactions were found to influence mAb production. Using the proteomics data of CHO cell lines producing another similar mAb in our lab [21], the enzyme expression levels of these metabolic reactions in high production cell line were compared to low production state. The predicted changes in the rates of these reactions in the FVSEOF list were in accordance with the enzyme expression changes (Supporting information, S0 file). For example, three enzymes of vitamin B6 pathway (namely, pyridoxamine kinase, pyridoxal kinase, and pyridoxine kinase) were overexpressed according to the proteome data and also their corresponding reactions were predicted to have increased rates in the FVSEOF list.

Experimental validation of the computational results, *i.e.*, adding each of these 15 metabolites to cell culture feeds, by changing one factor at a time could be laborious and time-consuming. In other words, considering the presence or absence of each of these 15 metabolites in cell culture feeds, one would need 2^15^=32768 sets of experiments, in a full factorial mode, to elucidate the exact effect of each feeds on CHO cells. Therefore, a DoE method has been used in this study to have an initial screening of the feeds and finding the most effective metabolites [22]. Plackett-Burman (PB) design method is one of the most widely used DoE methods [23], which has been usually used for screening the factors affecting an experiment [24,25]. PB method is based on the basic factorial design of experiments, which helps in measuring the effect of each variable component on an assumed response while holding the levels of the other components fixed, as well as when changing the levels of two or more components simultaneously [26]. The number of experiments in a full factorial design is increased exponentially by increasing the number of components, while PB design is simple and needs very few experiments to screen the effects of components. However, the PB method neglects the interactions of components and their synergistic effects, and only considers the main effects of them. In this study, a 20-run PB designed experiment was used to explore the effects of 15 metabolites on mAb production in CHO cells.

Statistical analysis of the results of PB design revealed that two metabolites, namely, arachidonate and threonine, can significantly improve mAb production in CHO cells. In order to find the most effective levels of concentrations of these two metabolites or increasing mAb production, response surface methodology (RSM) has been used. RSM is intended to explore the relationships between one or more response and several explanatory variables [27]. It was introduced by George E. P. Box and K. B. Wilson in 1951. They suggested using a second-degree polynomial model to approximate the relationship between variables and responses. In contrast to the PB method, the interaction among variables can be determined by statistical techniques [28]. Central composite design (CCD) is one of the response surface methods that aims to fit a model by least squares technique [29]. Adequacy of the proposed model is then revealed using the diagnostic checking tests provided by analysis of variance (ANOVA). Here, we used CCD to find the concentrations of threonine and arachidonate that can increase the mAb expression more effectively.

To start the experimental validation tests, some pre-experiments were performed to determine the initial concentrations of 15 metabolites in CHO cell culture medium (proCHO5), using high-performance liquid chromatography (HPLC) and gas chromatography-mass spectrometry (GC-MS). The initial concentrations of 7 amino acids and vitamin B1 in the CHO cell culture medium (proCHO5) were successfully determined (Supporting information, S1 file). It is assumed that the remaining 7 metabolites are not present in the cell culture medium.

To start experimental validation of the cell culture feeds using DoE method, we defined a concentration for each of these 15 metabolites in the feeds, which is obviously higher than the initial concentrations of metabolites in cell culture medium (proCHO5). The concentrations of amino acids were selected according to the available commercial feeds that were previously analyzed in our lab [30]. The concentrations of vitamin B1 and B6 were selected according to some available patents, like US5316938A, US9321996, and US4816401. The literature data regarding vitamin A and 5 other metabolites (thymidine, deoxy-cytidine, 3-methyl-oxobutyrate, deoxy-guanosine, and arachidonate) were not found. Therefore, we performed some pre-experiments to select suitable concentrations for these 6 metabolites (Supporting information, S2 file). The results of using DoE methods are as follow.

### Plackett-Burman design

The metabolic modeling predicted that increasing consumption rates of 15 metabolites might enhance the mAb production *in vitro*. To validate this prediction, these 15 metabolites were added to the feed solutions of CHO cells, using a matrix that was designed by the Plackett-Burman (PB) method (Figure 1, left panel). According to the PB design matrix, a total of 20 trials were performed at various combinations of ‘high’ (+) and ‘low’ (-) levels of concentrations of different metabolites. The low levels were the amount of metabolites in the CHO cell culture medium (proCHO5), and high values were determined according to the literature and some pre-experiments (Supporting information, S1 and S2 files).

**Figure 1.**
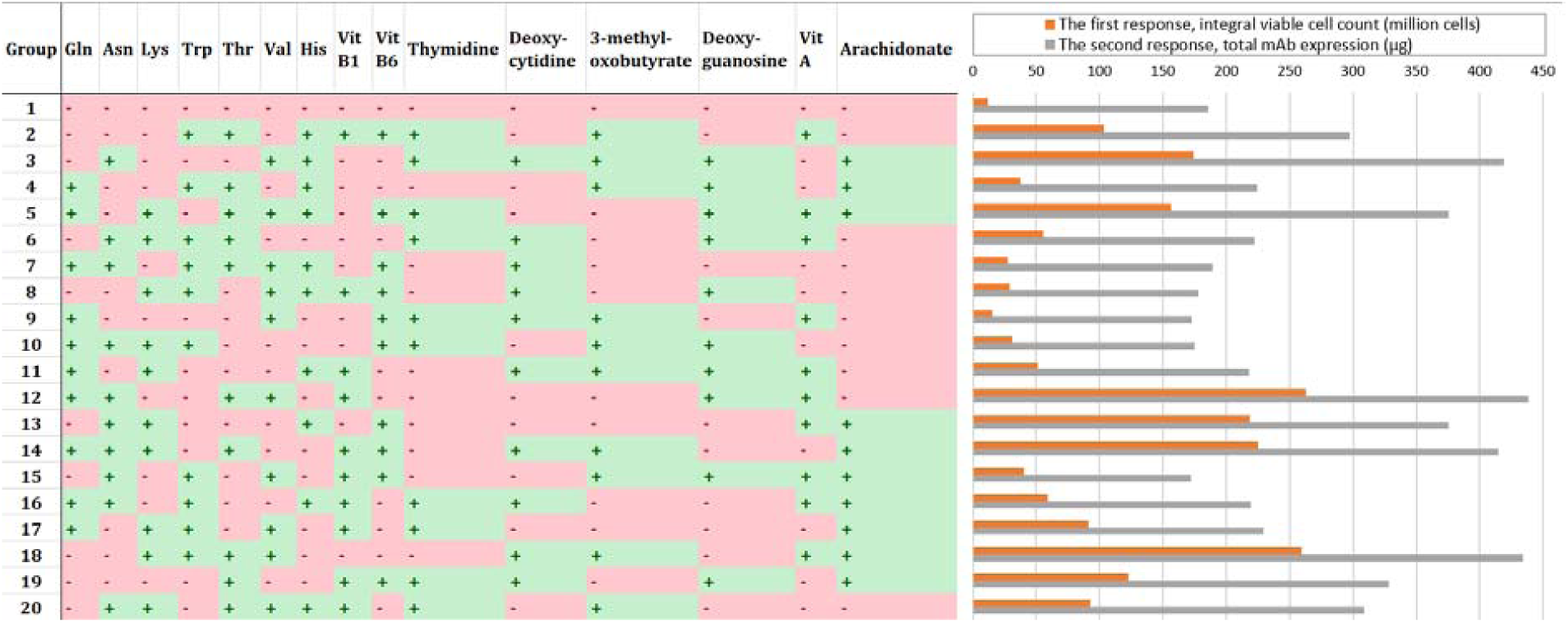
In the left part of the figure, the composition of CHO cell culture feeds in Plackett–Burman (PB) design matrix have been shown. Twenty groups of feed solutions were designed at various combinations of ‘high’ (+) and ‘low’ (-) levels of concentrations of 15 different metabolites. The exact amounts of the concentrations are included in Supporting information 1 and 2 (S1 and S2 files). Group 1 was considered as the negative control, because it does not contain any of the 15 metabolites as supplements and therefore, only proCHO5 medium was added to the cells as feed 1. Gln: glutamine, Asp: asparagine, Lys: lysine, Trp: tryptophan, Thr: threonine, Val: valine, His: histidine, Vit B1: vitamin B1 or thiamine, Vit B6: vitamin B6 or pyridoxine, Vit A: vitamin A or retinol. In the right part of the picture, the responses of Plackett–Burman design of experiments have been shown in a bar chart. The first response is integral viable cell count (IVCC), which is presented in million cells for each group. The second response is total mAb production.

The number of viable and dead cells were counted on each day for all groups. The detailed changes in cell viabilities and integral viable cell count (IVCC) have been shown in Figure S1 in Supporting information, S3 file. In order to have a representation of changes in the IVCC during cell culture, Hill equation was fitted to changes in the IVCC of CHO cells (Figure 2). In this equation, IVCC_max_ is the maximum amount of IVCCs during cell culture, which equals the IVCC in the harvesting day. D represents the times that CHO cells are in cell culture medium (in days). D_50_ equals the day in which the cells reach the half amount of their maximum IVCC, which has been shown in Figure 2. The groups that have bigger D_50_ values, have longer cell culture duration and therefore, better responses are expected in that group (e.g., group 12). The details about fitting are included in Supporting information, S3 file. The IVCC of the harvesting day and the total mAb expression of each group were used as responses in PB design experiments, which have been shown in a bar chart in Figure 1. According to PB matrix, group 1 did not contain any of the 15 supplemented metabolites and therefore, this group was considered as the control group. The highest total mAb production belongs to group 12, with 438 µg, which shows more than a two-fold increase in mAb production compared to control (186 µg). The detailed statistical analysis of the responses are presented in supporting information, S3 file. In brief, the results showed that two metabolites, namely, arachidonate and threonine, are the most influential supplemented metabolites that showed significant improvements in mAb production. Therefore, these two metabolites were chosen for further analysis using the RSM method.

**Figure 2.**
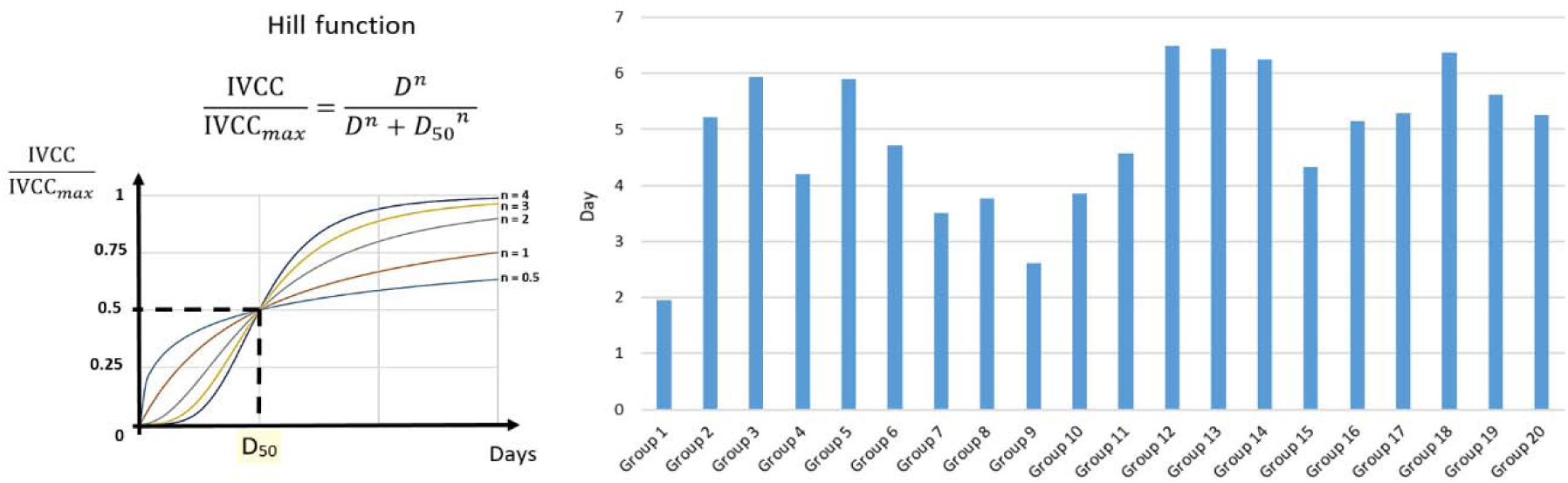
Hill equation and a schematic representation of it have been shown in the left side of the figure. In Hill equation, IVCC_max_ is the maximum amount of IVCCs during cell culture, which equals the IVCC in the harvesting day. D represents the times that CHO cells are in cell culture medium (in days). D_50_ equals the day in which the cells reach the half amount of their maximum IVCC. For each group, D_50_ values have been shown in the right side of the picture. Details about these data are included in Supporting information, S3 file.

### Central composite design

Arachidonate and threonine, were the most positive influential supplements that could significantly improve mAb production in PB designed experiments. To find the most effective concentrations of these two metabolites, central composite design (CCD) was used. Five levels of concentrations for each of the two metabolites were tested in 11 trials in CCD (Figure 3, left panel). The concentrations of metabolites in ‘0.00’ levels in CCD were equal to ‘+1.00’ or high levels in PB design. The details about CCD are included in Supporting information 4 (S4 file).

**Figure 3.**
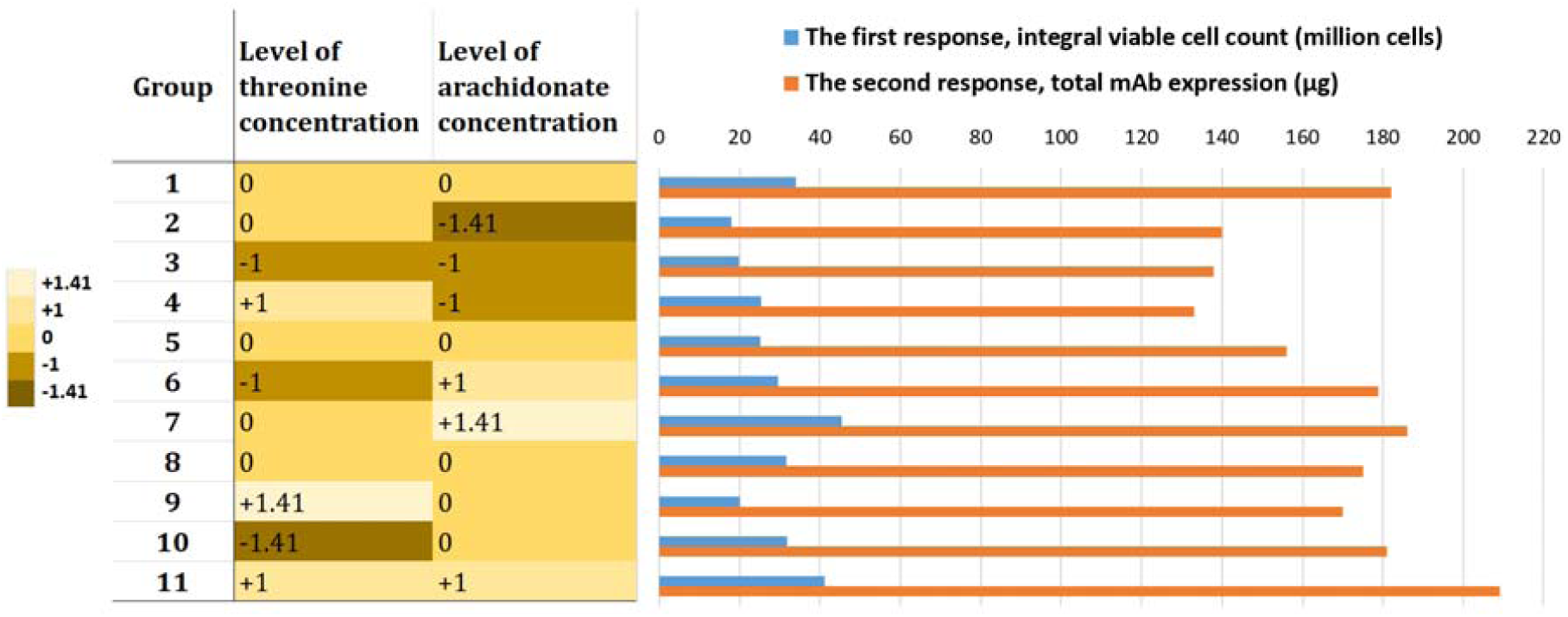
In the left part of the figure, the composition of CHO cell culture feeds in central composite design (CCD) matrix. Eleven groups of feed solutions were designed at various combinations of ‘+1.00’, ‘-1.00’, ‘0.00’, ‘+1.41’, and ‘-1.41’ levels of arachidonate and threonine concentrations. The details about CCD are included in Supporting information 4 (S4 file). In the right part of the picture, the responses of CCD have been shown in a bar chart. The integral viable cell count (IVCC) of each group is calculated in million cells. The second response is the total mAb expression, which has been determined in µg, using HPLC.

The number of viable and dead cells were counted on each day for all groups. Changes in cell viabilities and integral viable cell count (IVCC) have been shown in Figure S2 in Supporting information, S4 file. The IVCC of the harvesting day and the total mAb expression of each group were used as responses in CCD experiments, which have been shown in a bar chart in Figure 3. The same as PB design, Hill equation was fitted to changes in the IVCC of CHO cells (Figure 4). The details about fitting are included in Supporting information, S4 file. The 11 groups in CCD design contained different concentrations of threonine and arachidonate (Figure 3, left panel). We add a control group to CCD experiments, where the feed solution was only 0.5 ml of basal CHO culture medium (proCHO5) with no additional threonine and arachidonate. All other conditions for control group were the same as other groups. In the control group, the IVCC was 14.437 million cells, and total mAb expression was 83 µg. Therefore, about 2.5 fold increase in total mAb expression was seen in group 11 compared to control. The detailed statistical analysis of the responses are represented in supporting information, S4 file.

**Figure 4.**
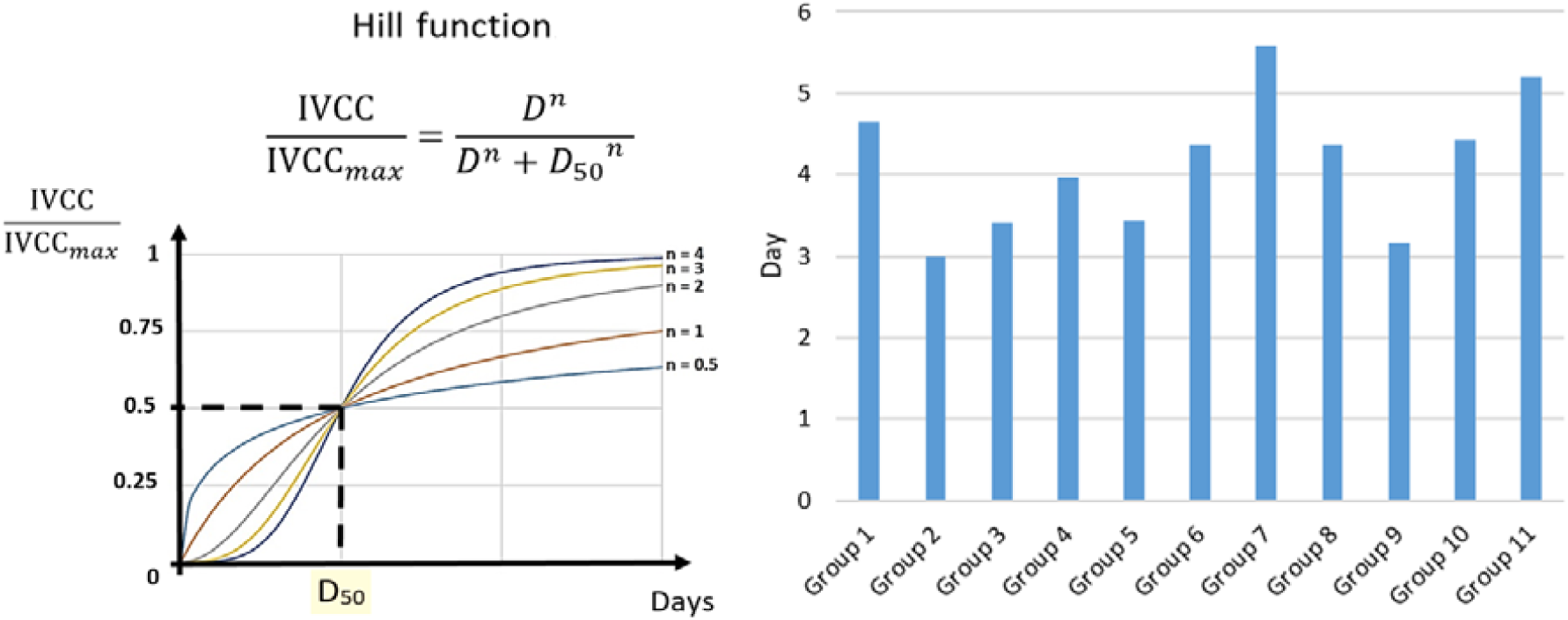
Hill equation and a schematic representation of it have been shown in the left side of the figure. In Hill equation, IVCC_max_ is the maximum amount of IVCCs during cell culture, which equals the IVCC in the harvesting day. D represents the times that CHO cells are in cell culture medium (in days). D_50_ equals the day in which the cells reach the half amount of their maximum IVCC. For each group of the CCD experiments, D_50_ values have been shown in the right side of the picture. Details about these data are included in Supporting information, S3 file.”

The 3-dimensional representations of the interaction of IVCC and total mAb expression with threonine and arachidonate concentration levels have been shown in Figure 5. It has been shown that both IVCC and mAb expression are directly related to the concentrations of threonine and arachidonate, and these two metabolites are positively interacting with each other in the studied range of our analysis.

**Figure 5.**
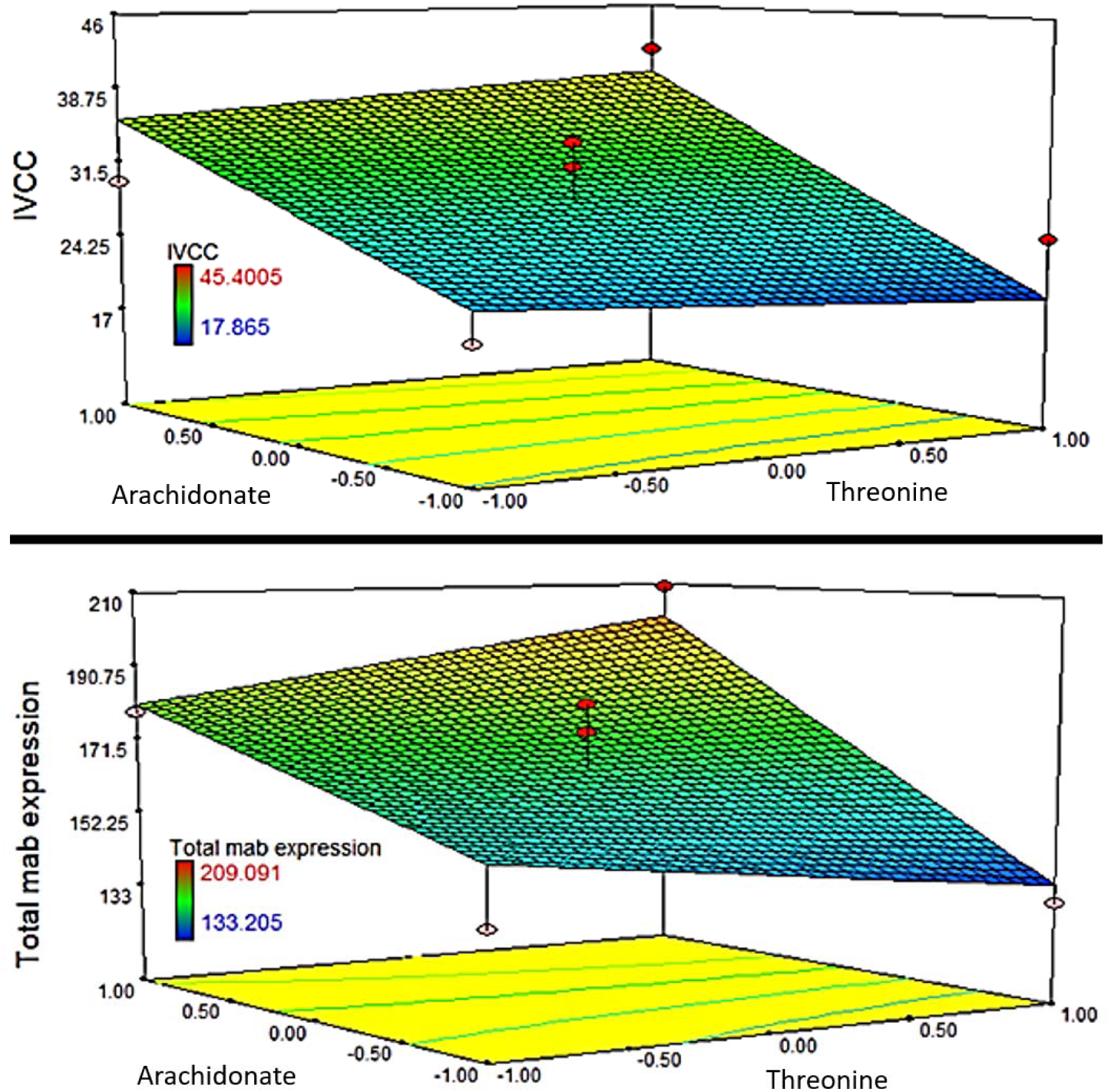
Representation of 3D response surface for integral viable cell count (IVCC) and total mAb expression. This response surface shows the interaction of metabolite A (threonine, on ‘x’ axis) and metabolite B (arachidonate, on ‘y’ axis) on IVCC (on ‘z’ axis, upper part of figure) and total mAb expression (on ‘z’ axis, lower part of figure). According to the statistical analysis in supporting information S4 file, IVCC = 29.29 + 0.08*A + 8.07* B + 1.52* AB and total mAb expression = 168.14 +1.12*A +22.65*B +8.84*AB. This means that both responses are directionally related to the concentrations of the two metabolites, and especially arachidonate. In addition, threonine and arachidonate are positively interacting with each other for increasing the responses.

## Discussion and Conclusions

In this study, a new approach for designing the compositions of feeds for increasing mAb production in CHO cell culture have been proposed and experimentally validated. To the best of our knowledge, this is the first time that a GEM of CHO cell line has been used for cell culture medium or feeding strategy design. Using the metabolites predicted by metabolic network modeling as culture supplements in feed solutions, about a two-fold increase in total mAb expression was achieved. According to the results, arachidonate was the most effective supplement in CHO cell culture feed for increasing mAb production. Arachidonate is one of the essential fatty acids [31]. It has been shown that dietary supplementation with arachidonate can improve cellular health and growth [32]. The major precursors of prostaglandins, prostacyclin, thromboxane, and eicosanoids have to be produced in the metabolism of arachidonate [33,34]. The positive effects of these arachidonate metabolites on the growth of Caco-2 cell line have been studied [35]. It has been shown that arachidonate can stimulate the proliferation of fetal mouse brain cells [36], human colon carcinoma cells [37], and human breast cancer cell lines [38]. Arachidonate can be synthesized from linoleic acid (another essential fatty acid). However, some studies indicated that some of the cultured cells, including CHO cells, are unable to synthesize arachidonate from linoleate, as CHO cells lacked enzyme activity for the desaturation, which is the first and usually rate-limiting step of the desaturation-elongation sequence [39]. Totally, most cells can synthesize fatty acids *in vitro*. However, *de novo* synthesis is usually relatively low and cultured cells tend to rely on exogenous free fatty acids. In conclusion, according to the mentioned literature, cell culture supplementation with arachidonate can impose positive effects on cell growth and proliferation. These positive effects are consistent with the current study and could be a validation of our results.

We note that there are some limitations in metabolic modeling. Even in the most complete metabolic models that have been reconstructed till now, some reactions may be missing, according to a lack of knowledge in cell biology or some errors made in the process of reconstruction of the model. In addition, a metabolic model only considers the biochemical reactions occurred in a cell, while a metabolite may also have a role in transcriptional or signaling pathways. In other words, increasing the consumption of a metabolite may increase the rate of a biochemical reaction in favor of mAb production, while this metabolite may also take part in a signaling pathway and cause up-regulation of another pathway that is against mAb production or against cell viability. Integration of signaling and metabolic networks may be a solution for these limitations [40]. However, there has been not any reliable signaling network for CHO cells until the writing of this study. Another limitation in metabolic modeling is choosing the right algorithm. As already mentioned, several strain design algorithms were tested in the first step of this study for analyzing the metabolic model of CHO cells, like optForce [41]. However, because of the relatively big size of the metabolic model of CHO cells, and also operational limitations of computational calculations, only FVSEOF algorithm came to reliable results.

In conclusion, our approach for fermentation process optimization, in which a metabolic modeling study was used upstream of the DoE methods as a means to reduce the number of variables in cell culture medium optimization, is suggested to be used for other cells and meabolic models.

## List of abbreviations

CHO: Chinese hamster ovary
mAbs: monoclonal antibodies
GEMs: genome-scale metabolic network models
GPR: gene-protein-reaction
FVSEOF: flux variability scanning based on enforced objective flux
DoE: design of experiment
PB: Plackett-Burman
RSM: response surface methodology
CCD: central composite design

## Acknowledgements

We have to kindly thank Samira Ahmadi and Radin Biotech company, Iran, for gifting the CHO cell line to be used in our study.

## Authors’ contributions

H.F. and S.-A.M. designed the computational studies. N.E.L. was involved in computational modeling of CHO cells metabolism. H.F., B.V., F.T., and F.M. designed the lab experiments. H.F. wrote the main manuscript. N.E.L., B.V., and S.-A.M. reviewed the manuscript. All authors read and approved the final manuscript.

## Data availability

All data generated or analyzed during this study are included in this published article and its supplementary information files. The metabolic model of CHO cells is publicly available in the supporting information of the original article (Hefzi et. al., Cell Systems, 2016), which has been cited in our article. The cell line which has been used in our study is available in Radin Biotech company of Iran.

## Compliance with ethical standards

### Human and animal rights statement

This article does not contain any studies with human participants or animals performed by any of the authors.

### Informed consent

Not applicable

## Supporting information

**S0 file:** Details about FVSEOF output.

**S1 file:** Determining initial concentrations of the 15 metabolic in CHO cell culture basal medium (proCHO5).

**S2 file:** Experimental pre-tests to choose concentrations for the 15 metabolites in feed solutions

**S3 file:** Detailed statistical analysis of the results of Plackett-Burman designed experiments.

**S4 file:** Detailed statistical analysis of the results of central composite designed experiments.

