## Supplementary file S0 for "A metabolic network-based approach for developing feeding strategies for CHO cells to increase monoclonal antibody production"

* Corresponding authors

**S0 file:**

**Details about FVSEOF output**

As mentioned in the manuscript, we used FVSEOF algorithm [1] to find the reactions that can influence mAb production. In other words, using the FVSEOF method and the metabolic network model of CHO cells, the effects of an enforced theoretical increase in mAb production on the rates of all other metabolic reactions of CHO cells were modeled. FVSEOF generated a list of more than 400 reactions which includes the metabolic reactions that have altered rates during the theoretical increase in mAb production. These altered rates were found by monitoring the changes in the flux bounds of reactions. If the direction of flux changes of a predicted reaction was in parallel with the enforced changes in mAb production, that predicted reaction was a candidate for increasing its rate *in vitro* to influence mAb production. In our study, we explored the list of candidates generated by FVSEOF to find the exchange reactions. We assumed that an increase in the rate of an exchange reaction can occurred by increasing the concentration of that exchanged metabolite in cell culture medium. Here, we report the directions of changes in the flux bounds of these candidate exchange reactions in a five step enforced theoretical increase in mAb production (Table S1). For each reaction, the average of minimum and maximum possible flux boundaries has been included in the table to show the trend of changes.

In addition, we found that some of the reactions in FVSEOF output list were reported to have a fold change in their catalyzing enzymes in low producing cell lines compared to high producing ones [2]. We have also included these data in the Table S1.

As shown in the table, during the enforced increase in mAb production from step 1 to step 5, the exchange fluxes of following 13 metabolites have been increased: glutamine, asparagine, lysine, tryptophan, threonine, valine, histidine, vitamin B1, thymidine, deoxy-cytidine, 3-methyl-oxobutyrate, deoxy-guanosine, and arachidonate. It has to be noted that consumption of a metabolite is modeled with a negative exchange flux of that metabolite in metabolic networks. In case of vitamin A, a decrease in the rate of its exchange toward extracellular space have been shown. Therefore, it was assumed that the CHO cell needs vitamin A for increasing mAb production. The increasing fluxes of pyridoxamine, pyridoxal, and pyridoxine kinase, which were in accordance with the fold changes in the enzymes of them, showed that vitamin B6 (pyridoxine) can be a good supplement for cell culture for increasing mAb production. Therefore, we used these 15 metabolites (13+1+1) in cell culture feeds. In Table S1, only one of the enzymes (Acetyl-CoA C-acetyltransferase) had decreased flux changes while the enzyme of that reaction was shown to have a two-fold change. This enzyme is suggested for further studies.

**Table S1**. FVSEOF results. In each steps of FVEOF, the average of minimum and maximum possible flux boundaries for each reaction have been shown.

| Rxn Name | Average of flux bounds in step 1 of FVSEOF | Average of flux bounds in step 2 of FVSEOF | Average of flux bounds in step 3 of FVSEOF | Average of flux bounds in step 4 of FVSEOF | Average of flux bounds in step 5 of FVSEOF | Enzyme ID | Fold change in proteome data |
| --- | --- | --- | --- | --- | --- | --- | --- |
| Biomass production | 281.496 | 250.7312 | 219.9663 | 189.2014 | 155.8537 | - | - |
| mAb production | 0.23 | 0.46 | 0.69 | 0.92 | 1.15 | - | - |
| Glutamine exchange | -570.897 | -572.078 | -573.26 | -574.441 | -575.046 | - | - |
| Asparagine exchange | -543.148 | -545.066 | -546.984 | -548.902 | -550.479 | - | - |
| Lysine exchange | -79.0938 | -81.6756 | -84.2574 | -86.8393 | -88.7882 | - | - |
| Tryptophan exchange | -507.239 | -509.51 | -511.78 | -514.051 | -516.28 | - | - |
| Threonine exchange | -557.169 | -563.933 | -570.697 | -577.461 | -583.808 | - | - |
| Valine exchange | -71.2053 | -78.7314 | -86.2576 | -93.7837 | -100.783 | - | - |
| Histidine exchange | -20.5699 | -22.1489 | -23.7278 | -25.3068 | -26.7287 | - | - |
| Vit B1 exchange | -161.33 | -161.913 | -162.496 | -163.08 | -163.712 | - | - |
| Thymidine exchange | -163.211 | -163.588 | -163.966 | -164.344 | -164.753 | - | - |
| Deoxy-cytidine exchange | -162.653 | -163.092 | -163.53 | -163.969 | -164.444 | - | - |
| 3-methyl-oxobutyrate exchange | -71.2053 | -78.7314 | -86.2576 | -93.7837 | -100.783 | - | - |
| Deoxy-guanosine exchange | 0 | 0 | 0 | 0 | -0.73266 | - | - |
| Arachidonate exchange | -437.5 | -437.5 | -437.5 | -437.5 | -668.834 | - | - |
| Vitamin A exchange | 16.01087 | 14.26102 | 12.51117 | 10.76132 | 8.864579 | - | - |
| Pyridoxamine kinase | 483.9891 | 485.739 | 487.4888 | 489.2387 | 491.1354 | PYDAMK | 1.8 |
| Pyridoxal kinase | 483.9891 | 485.739 | 487.4888 | 489.2387 | 491.1354 | PYDXK | 1.8 |
| Pyridoxine kinase | 483.9891 | 485.739 | 487.4888 | 489.2387 | 491.1354 | PYDXNK | 1.8 |
| Nucleoside-diphosphate kinase (ATP:dUDP) | -330.297 | -330.629 | -330.96 | -331.292 | -331.652 | NDPK6 | 1.8 |
| RE0383 | 44.13765 | 188.1521 | 335.6773 | 483.2024 | 500 | RE0383C | 1.7 |
| Glucose 6-phosphate dehydrogenase | 455.8624 | 455.8386 | 455.8148 | 455.7911 | 456.1322 | G6PDH2r | 1.8 |
| Hydroxymethylglutaryl CoA synthase (ir) | 500 | 500.282 | 501.3931 | 502.5041 | 504.5999 | HMGCOASim | 2.3 |
| Acetyl-CoA C-acetyltransferase | 166.36 | 166.0533 | 165.7467 | 165.44 | 165.1333 | ACACT1r | 2 |
