## Supplementary file S1 for "A metabolic network-based approach for developing feeding strategies for CHO cells to increase monoclonal antibody production"

* Corresponding authors

**S1 file:**

**Determining initial concentrations of the 15 metabolic in CHO cell culture basal medium (proCHO5)**

**1. Introduction**

As mentioned in the text, the results of analyzing CHO metabolic model with the FVSEOF algorithm [1] predicted that increasing the consumption of 7 amino acids (glutamine, asparagine, lysine, tryptophan, threonine, valine, and histidine), 3 vitamins (vitamin A, B1 and B6), and 5 other metabolites (thymidine, deoxy-cytidine, 3-methyl-oxobutyrate, deoxy-guanosine, and arachidonate) in CHO cells can increase *in vitro* production of monoclonal antibody (mAb). To validate the computational predictions, a Plackett-Burman (PB) method was used to design the experiment of adding these 15 metabolites to cell culture medium as feeds. The initial concentrations of these 15 metabolites in CHO cell culture basal medium (proCHO5) have to be determined firstly. These concentrations were (-1) levels of the metabolite in PB design. Here, some pre-experiments were performed for this matter, using high-performance liquid chromatography (HPLC) and gas chromatography-mass spectrometry (GC-MS).

**2. Materials and Methods**

*2.1. HPLC*

*2.1.1. Amino acids*

In the first part, the concentrations of amino acids in CHO cell culture basal medium (proCHO5) were determined. For this purpose, amino acids were derivatized using OPA (Pickering Laboratories, CA, USA) and separated by HPLC, using a C18 column with a florescence detector [2]. A standard solution of amino acids with defined concentrations was also used in HPLC with the same conditions. The concentration of each amino acid is relative to its peak area in the chromatogram. Therefore, the unknown concentrations in proCHO5 were calculated by comparing the peak area in two chromatograms.

It has to be noted that glutamine and histidine peaks have the same retention times, so we cannot determine the concentration of each of them (Figure 1). The same happens for tryptophan and methionine. Therefore, the concentrations of these 4 amino acids cannot be determined with the previous run of HPLC. To solve this problem for glutamine and histidine peaks, an acid hydrolysis treatment was performed for the solutions before running them in HPLC. In such condition, the glutamine content of the solution is converted to glutamic acid. Therefore, the peaks are separated and the concentration of histidine can be calculated. In addition, the total amount of glutamic acid in the acidic solution is calculated, and by reducing the concentration of glutamic acid (which is previously determined in non-acid hydrolyzed condition) from the total concentration of glutamic acid (which is determined in the acidic solution), the concentration of glutamine in proCHO5 will be calculated easily. Therefore, in the second run of HPLC in the acid hydrolyzed condition, the concentrations of histidine and glutamine were calculated. However, the concentrations of tryptophan and methionine remained unknown (see the 2.1.2).

*2.1.2. Vitamins*

To determine the concentrations of water- and fat-soluble vitamins in proCHO medium simultaneously, we used previously described methods [3]. In brief, a C18-A (250mm × 4.6mm, 5µm particle size) was used for HPLC with a UV detector, at 280 nm wavelength. The mobile phase consists of 0.010% triﬂuoroacetic acid (solvent A) and methanol (solvent B), at the ﬂow rate 0.7 ml/min. A linear gradient proﬁle (A:B) started at 95:5 and was constant for the ﬁrst 4 min, then it linearly decreased to 2:98 during the next 6 min and was constant for 20 min, and ﬁnally linearly increased up to 95:5 for the last 5 min of the separation.

Some amino acids, namely tryptophan, tyrosine, and phenylalanine, absorb UV light at 280 nm wavelength. The standard solutions of these amino acids were also used in the current HPLC method. Therefore, the concentration of tryptophan was successfully determined using this method. In the first part, total concentration of tryptophan and methionine had been calculated (see 2.1.1). Therefore, the concentration of methionine was calculated easily.

*2.2. GC-Mass analysis*

To determine the concentrations of other metabolites in proCHO5 medium, GC-Mass analysis was used. The samples that are analyzed by GC-Mass should not be in aqueous solution. Therefore, 1 ml of proCHO5 medium was evaporated in room temperature using a concentrator. Then, the contents were dissolved in 1 ml of methanol. The solution was filtered and used in GC-Mass analysis.

*2.2.1. Temperature Program:*

Initial temperature (50 ºC), initial time (1 min), program rate (10 ºC/min), final temperature (250 ºC), final time (10 min), split ratio (25 ml/min), flow rate (1 ml/min).

*2.2.2. Instrument Specifications:*

The manufacturer company was Agilent Technologies, with 7890A GC System. The mass selective detector was 5975C VL MSD (Triple-Axis Detector). The Ion source was Electron Impact (EI) 70eV. A quadrupole analyzer and Rtx 5 MS column with 30 m length, 0.250 mm I.D. and a film thickness of 25 μm was used. Data were processed by Agilent MSD Chemstation (Rev E.02.02.1413). Injection port temperature was 250 ºC, detector temperature was 230 ºC. The carrier gas was He (99.999%). The sample volume was 1 μL.

**3. Results** **and** **Discussion:**

*3.1. HPLC analysis*

As mentioned in the methods, three HPLC runs were performed (Figure 1-3). The calculated concentrations of amino acids and vitamins have been shown in Table 1. **These concentrations were used as (-1) levels in Plackett-Burman design of experiment.**

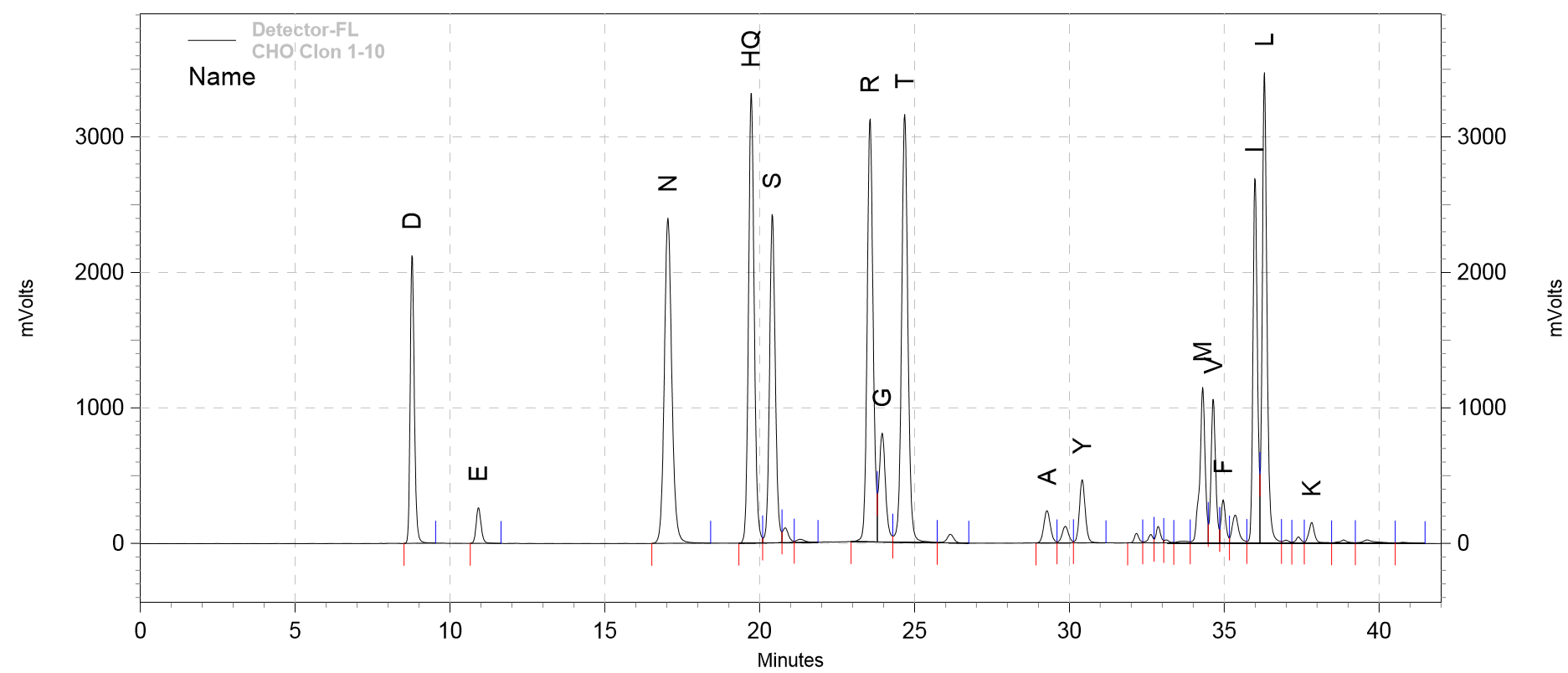

**Figure 1**. HPLC chromatogram of proCHO5 medium in the first run. The peak denoted by ‘HQ’ belongs to histidine and glutamine, and the peak denoted by ‘M’ belongs to methionine and tryptophan. Other abbreviations are as follow: D (Asp), E (Glu), N (Asn), S (Ser), R (Arg), G (Gly), T (Thr), A (Ala), Y (Tyr), V (Val), F (Phe), I (Ile), L (Leu), K (Lys).

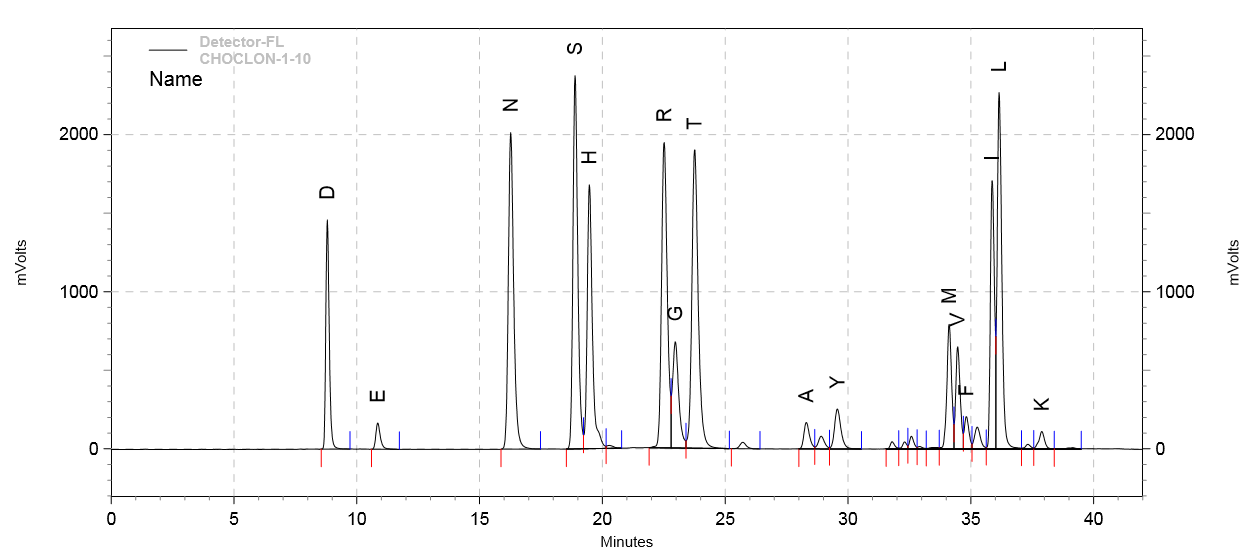

**Figure 2**. HPLC chromatogram of proCHO5 medium in the second run (acid hydrolyzed condition). The peak denoted by ‘M’ belongs to methionine and tryptophan. Other abbreviations are as follow: D (Asp), E (Glu+Gln), N (Asn), S (Ser), H (His), R (Arg), G (Gly), T (Thr), A (Ala), Y (Tyr), V (Val), F (Phe), I (Ile), L (Leu), K (Lys).

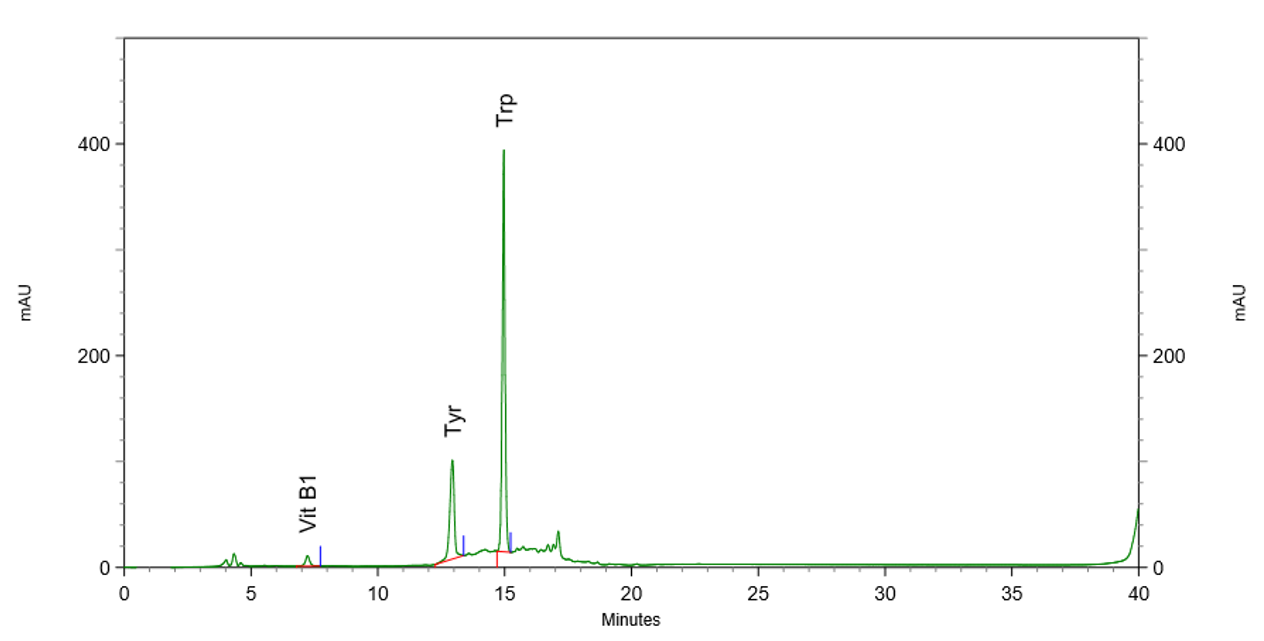

**Figure 3**. HPLC chromatogram of proCHO5 medium in the third run (considering vitamins and tryptophan). Phenylalanine was not detectable using this method. Additionally, the retention time of ‘vitamin A’ is 21.5 minutes, and ‘vitamin B6’ is 14 minutes, but we did not detect these vitamins in proCHO5 medium.

**Table 1**. The concentrations of proCHO5 components, calculated by HPLC.

| **Names of the components** | **Concentrations in proCHO5 medium (mg/ml)** |
| --- | --- |
| D (Asp) | 0.70293431 |
| E (Glu) | 0.45430301 |
| N (Asn) | 0.74894423 |
| S (Ser) | 0.69520648 |
| R (Arg) | 0.66501214 |
| G (Gly) | 0.25774407 |
| T (Thr) | 0.75726153 |
| A (Ala) | 0.04924609 |
| Y (Tyr) | 0.16453328 |
| V (Val) | 0.13881608 |
| F (Phe) | 0.07688753 |
| I (Ile) | 0.38322629 |
| L (Leu) | 0.58430895 |
| K (Lys) | 0.30807529 |
| Q (Gln) | 0.08200175 |
| H (His) | 0.38801192 |
| W (Trp) | 0.17016065 |
| M (Met) | 0.14156505 |
| Vitamin B1 | 0.00832783 |
| Vitamin B6 | 0 |
| Vitamin A | 0 |

*3.2. GC-Mass analysis*

As we mentioned in the introduction, the modeling results predicted that increasing the consumption of 15 metabolites might increase mAb production in CHO cells. The concentrations of 10 metabolites (7 amino acids and 3 vitamins) were determined by HPLC. In order to determine the concentrations of the remaining 5 metabolites (thymidine, deoxy-cytidine, 3-methyl-oxobutyrate, deoxy-guanosine, and arachidonate) in proCHO5 medium, GC-MS was used. According to the results (Figure 4 and Table 2), any of these 5 metabolites were detected in proCHO5 medium. Therefore, the concentrations of these metabolites were assumed to be zero in proCHO5 medium. In other words, the **(-1) levels in Plackett-Burman design of experiment for these 5 metabolites were zero.**

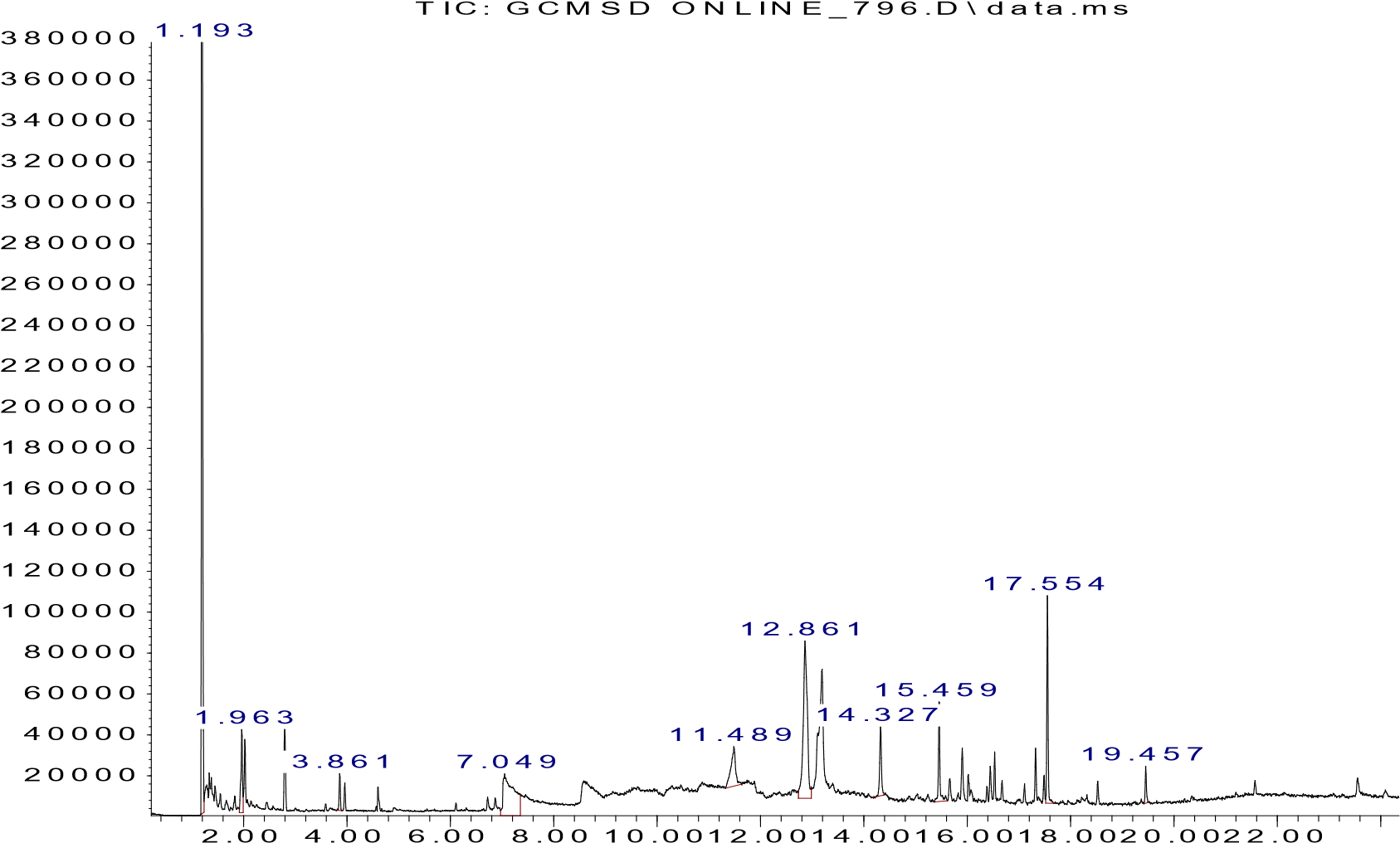

**Figure 4**. GC histogram.

**Table 2**. GC-Mass results. The numbers in the 4^th^ column (A%) can be accounted as w/w%.

| **Compound number** | **Compound Name** | **RT (min)** | **Area (Ab*s)** | **A%** |
| --- | --- | --- | --- | --- |
| 1 | Carbon dioxide | 1.196 | 12399609 | 40.577 |
| 2 | 3-methyl-Butanal | 1.969 | 716371 | 2.344 |
| 3 | 2-methyl-Butanal | 2.028 | 484381 | 1.585 |
| 4 | 2,2-dimethoxybutane | 2.802 | 778144 | 2.546 |
| 5 | Glycine, methyl ester | 3.863 | 277546 | 0.908 |
| 6 | Triethylenediamine | 7.049 | 1588482 | 5.198 |
| 7 | Cyclodecanol | 11.492 | 1197600 | 3.919 |
| 8 | 2,6-di(t-butyl)-4-hydroxy-4-methyl-2,5cyclohexadien-1-one | 12.86 | 3729653 | 12.205 |
| 9 | pentadecane | 13.115 | 747554 | 2.446 |
| 10 | 10-Undecen-1-ol acetate | 13.191 | 2760442 | 9.033 |
| 11 | Hexadecane | 14.33 | 795371 | 2.603 |
| 12 | Heptadecane | 15.46 | 1052821 | 3.445 |
| 13 | 1H-Indene, 2,3-dihydro-1,1,3-trimethyl-3-phenyl | 15.901 | 499915 | 1.636 |
| 14 | 2H-Pyrido(2,1-b)(1,3)oxazinium | 16.454 | 319945 | 1.047 |
| 15 | Octadecane | 16.53 | 344424 | 1.127 |
| 16 | 9-Nonadecene | 17.329 | 438206 | 1.434 |
| 17 | 19-Formyloxy-6,11,12,14-tetrahydroxy-13-(2hydroxypropyl)-13-desisopropylabieta-5,8,11,13tetraen-7-one | 17.49 | 232415 | 0.761 |
| 18 | Nonadecane | 17.558 | 1656186 | 5.420 |
| 19 | 2-ethyl-2-methyl-Tridecanol | 18.526 | 248135 | 0.812 |
| 20 | Hexadecane | 19.461 | 291157 | 0.953 |

**4. Conclusion**

As mentioned, a PB design of experiment was used to validate the computational predictions made in our study. The initial concentrations of these 15 predicted metabolites in CHO cell culture basal medium (proCHO5) were used as (-1) levels. In the current supplementary file, these concentrations were determined and presented in Table 3.

**Table 3**. The final concentrations of the 15 metabolites that were used as (-1) level in Plackett-Burman design of experiment.

| Name of metabolite | Concentration (mg/ml) |
| --- | --- |
| Gln | 0.082 |
| Asn | 0.749 |
| Lys | 0.308 |
| Trp | 0.170 |
| Thr | 0.757 |
| Val | 0.139 |
| His | 0.388 |
| Vitamin B1 | 0.008 |
| Vitamin B6 | 0 |
| Thymidine | 0 |
| Deoxycytidine | 0 |
| Arachidonate | 0 |
| Deoxyguanosine | 0 |
| 3-methyl-oxobutyrate | 0 |
| Vitamin A | 0 |
