## Supplementary file S2 for "A metabolic network-based approach for developing feeding strategies for CHO cells to increase monoclonal antibody production"

* Corresponding authors

**S2 file:**

**Experimental pre-tests to choose concentrations for the 15 metabolites in feed solutions**

**1. Introduction**

As mentioned in the text, the results of analyzing the CHO metabolic model with the FVSEOF algorithm [1] predicted that increasing the consumption of 7 amino acids (glutamine, asparagine, lysine, tryptophan, threonine, valine, and histidine), 3 vitamins (vitamin A, B1 and B6), and 5 other metabolites (thymidine, deoxy-cytidine, 3-methyl-oxobutyrate, deoxy-guanosine, and arachidonate) in CHO cells can increase *in vitro* production of monoclonal antibody (mAb). To validate the computational predictions, these 15 metabolites have to be added to cell culture medium as feeds. The initial concentrations of these 15 metabolites in CHO cell culture basal medium (proCHO5) were determined in supplementary file 1 (S1 file). These concentrations were used as (-1) levels in Plackett-Burman design of experiment. The (+1) levels of concentrations for each of these 15 metabolites had to be defined to be used in cell culture feeds, which were obviously higher than the initial concentrations of metabolites in proCHO5. The concentrations of amino acids were selected according to the available commercial feeds that were previously analyzed in our lab [2]. The analysis method was the same as the method mentioned in S1 file. The concentrations of vitamin B1 and B6 were the highest amounts mentioned in the available patents, like US5316938A, US9321996, and US4816401. The literature data regarding vitamin A and 5 other metabolites (thymidine, deoxy-cytidine, 3-methyl-oxobutyrate, deoxy-guanosine, and arachidonate) were not found. Therefore, some pre-experiments have been performed here, to choose concentrations for the 15 metabolites in feed solutions. **The final concentrations of the 15 metabolites that were used as (+1) levels in Plackett-Burman design of experiment have been mentioned in Table 5.**

**2. Materials and Methods**

2.1. Cell culture

The CHO cell line, producing an anti α4β7 antibody, was gifted from PersisGene Company, Iran. The cells were cultivated in proCHO5 cell culture medium (purchased from Lonza AG, Verviers, Belgium), supplemented with 4 mM L-glutamine, 0.1% of anti-clumping agent, 1% of pluronic F68 and 1% of Pen-Strep (Gibco, Life Technology, USA).

CHO cells were cultured in 20 ml glass bottles (Duran Schott®) with a working volume of 5 ml, incubated at 37°C with a 5% solution of CO_2_, and agitated at 120 rpm. Each vessel was inoculated with a cell density of 5×10^5^ viable cells/ml, when the viability was more than 90%. The cells were cultivated in fed-batch mode and were supplemented with 0.5 mL of the feed solutions on day 3, 5, and 7.

Feed solutions includes different composition of 6 metabolites (vitamin A and the other 5 metabolites: thymidine, deoxy-cytidine, 3-methyl-oxobutyrate, deoxy-guanosine, and arachidonate) dissolved in proCHO5 cell culture medium. The vitamin A and other metabolites were purchased from Sigma-Aldrich Company (Germany).

Every day, 50 µL of cell culture medium were used for analyzing the viabilities of the cells. Cell viability was estimated using Trypan Blue assay, using a Neubauer cytometer. The cells were harvested when the viabilities dropped to less than 50%. In these pre-experiments, changes in the viabilities of the cells were used as the first reason to choose the best concentration for each metabolite. In addition, the integral viable cell count (IVCC) of the harvesting day was used as the second reason. IVCC was calculated by cumulative addition of viable cell counts (calculated in million cells) of each day of cell culture.

**3. Results:**

As mentioned, the concentrations of the amino acids and vitamins in the feed solutions were defined according to the literature data. However, any related literature data regarding the concentrations of vitamin A and the five remaining metabolites, i.e. thymidine, deoxycytidine, arachidonate, deoxyguanosine, and 3-methyl-oxobutyrate, in CHO cell culture for increasing cell proliferation or recombinant was not found.

*3.1. Thymidine, deoxycytidine, arachidonate, deoxyguanosine, and 3-methyl-oxobutyrate*

Three level of concentrations were chosen for each metabolite in the pre-experiments (Table 1).

**Table 1**. The concentrations of 5 metabolites that were used in the pre-experiments.

| **Name of metabolite** | **Thymidine** | **Deoxycytidine** | **Arachidonate** | **Deoxyguanosine** | **3-methyl-oxobutyrate** |
| --- | --- | --- | --- | --- | --- |
| **First level of concentration (mg/ml)** | 12 | 8 | 0.120 | 0.267 | 10 |
| **Second level of concentration (mg/ml)** | 6 | 4 | 0.040 | 0.027 | 5 |
| **Third level of concentration (mg/ml)** | 3 | 1 | 0.012 | 0.007 | 1 |

During the tests, cell count and viabilities were monitored on each day. The alterations of cell viabilities for each metabolites have been shown in Figure 1-5. Additionally, the integral viable cell count (IVCC) of the harvesting day was calculated by cumulative adding of cell densities, as shown in Table 2.

Thymidine

Viability (%)

| Time after cell culture | Control | First level | Second level | Third level |
| --- | --- | --- | --- | --- |
| Day 1 | 99 | 99 | 99 | 99 |
| Day 2 | 97.3 | 98.1 | 92.8 | 97.5 |
| Day 3 | 97 | 98.55 | 95.65 | 98.25 |
| Day 4 | 96.7 | 99 | 98.5 | 99 |
| Day 5 | 88.3 | 91.6 | 91.9 | 95.2 |
| Day 6 | 91.7 | 92.8 | 87.7 | 91.1 |
| Day 7 | 83.8 | 78.7 | 76.7 | 85.6 |
| Day 8 | 62.2 | 66.6 | 85.7 | 73.8 |

**Figure 1**. Changes in CHO cell viabilities during cell culture supplemented with thymidine. The concentrations of each level is detailed in Table 1.

Deoxycytidine

Viability (%)

| Time after cell culture | Control | First level | Second level | Third level |
| --- | --- | --- | --- | --- |
| Day 1 | 99 | 99 | 99 | 99 |
| Day 2 | 97.3 | 94.6 | 97.6 | 95.3 |
| Day 3 | 97 | 96.1 | 97.05 | 96.2 |
| Day 4 | 96.7 | 97.6 | 96.5 | 97.1 |
| Day 5 | 88.3 | 95 | 94.5 | 96.2 |
| Day 6 | 91.7 | 93.9 | 97.1 | 91.2 |
| Day 7 | 83.6 | 74.3 | 81.5 | 81 |
| Day 8 | 62.2 | 80.9 | 63.8 | 77.4 |

**Figure 2**. Changes in CHO cell viabilities during cell culture supplemented with deoxycytidine. The concentration of each level is found in Table 1.

Arachidonate

Viability (%)

| Time after cell culture | Control | First level | Second level | Third level |
| --- | --- | --- | --- | --- |
| Day 1 | 99 | 99 | 99 | 99 |
| Day 2 | 96.8 | 96.2 | 96.2 | 96 |
| Day 3 | 92.5 | 92.5 | 92.5 | 92.5 |
| Day 4 | 98.1 | 95.3 | 98.3 | 97.2 |
| Day 5 | 95.6 | 90.9 | 97.5 | 93.5 |
| Day 6 | 84 | 85 | 94.9 | 94.7 |
| Day 7 | 93.1 | 92.2 | 93.3 | 91.5 |
| Day 8 | 81.4 | 82.5 | 87.2 | 87 |

**Figure 3**. Changes in CHO cell viabilities during cell culture supplemented with arachidonate. The concentration for each level is detailed in Table 1.

Deoxyguanosine

Viability (%)

| Time after cell culture | Control | First level | Second level | Third level |
| --- | --- | --- | --- | --- |
| Day 1 | 82.69 | 82.69 | 82.69 | 82.69 |
| Day 2 | 83.03 | 82.68 | 89.02 | 93.86 |
| Day 3 | 94.26 | 84.78 | 95.89 | 95.78 |
| Day 4 | 89.14 | 83.19 | 96.45 | 86.38 |
| Day 5 | 57.78 | 72 | 88.03 | 81.86 |

**Figure 4**. Changes in CHO cell viabilities during cell culture supplemented with deoxyguanosine. The concentration for each level is detailed in Table 1.

3-methyl-oxobutyrate

Viability (%)

| Time after cell culture | Control | First level | Second level | Third level |
| --- | --- | --- | --- | --- |
| Day 1 | 95 | 95 | 95 | 95 |
| Day 2 | 89.65 | 90.35 | 93.2 | 93.2 |
| Day 3 | 84.3 | 85.7 | 91.4 | 91.4 |
| Day 4 | 90.2 | 93.3 | 89.9 | 96 |
| Day 5 | 91 | 91.8 | 88.3 | 94.2 |
| Day 6 | 65.3 | 89.1 | 92.7 | 89.3 |

**Figure 5**. Changes in CHO cell viabilities during cell culture supplemented with 3-methyl-oxobutyrate. The concentration for each level is detailed in Table 1.

**Table 2**. The integral viable cell count (IVCC) of the harvesting day for each group of metabolites, in three levels of concentrations, and the control of each group, calculated in million viable cells.

| **Name of metabolite** | **Thymidine** | **Deoxycytidine** | **Arachidonate** | **Deoxyguanosine** | **3-methyl-oxobutyrate** |
| --- | --- | --- | --- | --- | --- |
| **IVCC of control group** | 32.91 | 35.4 | 27.455 | 11.54 | 15.045 |
| **IVCC of the first level of concentration** | 33.905 | 37.705 | 30.925 | 3.89 | 17.82 |
| **IVCC of the second level of concentration** | 40.455 | 38.365 | 37.965 | 11.735 | 16.86 |
| **IVCC of the third level of concentration** | 41.735 | 42.415 | 35.425 | 14.08 | 16.73 |

*3.2. Vitamin A*

Vitamin A (powder) was not dissolved in cell culture medium (proCHO5). Therefore, according to [3], 1,4-dioxane was chosen as solvent. We have also tested ethanol as another solvent. The feed stocks of ‘Vitamin A’ groups were prepared by dissolving 0.1 mg of vitamin A powder in 10 µL of dioxane, 0.1 mg of vitamin A powder in 10 µL of ethanol, and 0.05 mg of vitamin A powder in 5 µL of ethanol and then adding proCHO5 medium to each solution to reach the total volume of 1 ml. The feed stock of ‘Control + Ethanol’ group was prepared by adding 10 µL of ethanol to 990 µL of proCHO5. The feed stock of the ‘Control+Dioxane’ group was prepared by adding 5 µL of dioxane to 995 µL of proCHO5. The feed stock of ‘Control’ group was only proCHO5.

During the tests, cell count and viabilities were monitored on each day. The alterations of cell viabilities for each group has been shown in Figure 6. Additionally, the integral viable cell count (IVCC) of the harvesting day was calculated by cumulative adding of cell counts, as shown in Table 3.

Vitamin A

Viability (%)

| Time after cell culture | Control | Control+Ethanol (1%) | Vit A in Dioxane (0.05 mg/ml) | Vit A in Ethanol (0.1 mg/ml) | Vit A in Ethanol (0.05 mg/ml) | Control+Dioxane (0.5%) |
| --- | --- | --- | --- | --- | --- | --- |
| Day 1 | 80 | 80 | 80 | 80 | 80 | 80 |
| Day 2 | 86.15 | 84.05 | 92.06 | 84.37 | 84.05 | 85.5 |
| Day 3 | 93.28 | 96.03 | 94.5 | 90.44 | 94.35 | 93 |
| Day 4 | 98 | 96.45 | 92.37 | 91.18 | 97.66 | 96 |
| Day 5 | 97.25 | 88.96 | 89.79 | 82.22 | 90.22 | 92 |
| Day 6 | 80.3 | 85.66 | 72.53 | 66 | 81.82 | 75.5 |
| Day 7 | 59.57 | 87.77 | 70.97 | 69.23 | 56.41 | 66.6 |

**Figure 6**. Changes in CHO cell viabilities during cell culture supplemented with vitamin A solutions and control groups. The feed stocks of ‘Vitamin A’ groups were prepared by dissolving 0.1 mg of vitamin A powder in 10 µL of dioxane, 0.1 mg of vitamin A powder in 10 µL of ethanol, and 0.05 mg of vitamin A powder in 5 µL of ethanol and then adding proCHO5 medium to each solutions to reach the total volume of 1 ml. The feed stock of the ‘Control+Ethanol’ group was prepared by adding 10 µL of ethanol to 990 µL of proCHO5. The feed stock of the ‘Control+Dioxane group was prepared by adding 5 µL of dioxane to 995 µL of proCHO5. The feed stock of the ‘Control’ group was only proCHO5.

**Table 3**. The integral viable cell count (IVCC) of the harvesting day for each group of vitamin A solutions, calculated in million viable cells.

| Group Name | IVCC |
| --- | --- |
| Control | 20.19 |
| Control+Ethanol (1%) | 29.86 |
| Control+Dioxane (0.5%) | 19.92 |
| Vit A in Dioxane (0.05 mg/ml) | 10.28 |
| Vit A in Ethanol (0.1 mg/ml) | 8.99 |
| Vit A in Ethanol (0.05 mg/ml) | 20.58 |

**4. Discussion:**

*4.1. Thymidine, deoxycytidine, arachidonate, deoxyguanosine, and 3-methyl-oxobutyrate*

According to Figure 1 and 3 and Table 2, in the case of thymidine and arachidonate, both of cell viability and IVCC have indicated consistent results and suggested that the best concentration for thymidine is the third one and for arachidonate is the second one. On the other hand, according to Figure 2 and 5 and Table 2, the results of cell viability and IVCC are not consistent for deoxycytidine and 3-methyl-oxobutyrate. In other words, the results of IVCC suggested that the third concentration for deoxycytidine and the first one for 3-methyl-oxobutyrate are the best concentrations, meanwhile, according to the results of cell viabilities, the second concentration is best for deoxycytidine and the third concentration is best for 3-methyl-oxobutyrate. Therefore, we used the average of the appropriate levels as ‘high’ levels for deoxycytidine and 3-methyl-oxobutyrate (Table 10). In case of deoxyguanosine, the second and third levels of the concentrations have shown positive effects on the viabilities of the cells. However, the IVCC of the third level is far more that the second level. Therefore, the third level, *i.e.*, 0.007 mg/ml is chosen for the future tests. In conclusion, the selected concentrations of the five metabolites have been shown in Table 4.

**Table 4**. The selected concentrations (mg/ml) of the five metabolites in cell culture feeds.

| Name of metabolite | Thymidine | Deoxycytidine | Arachidonate | Deoxyguanosine | 3-methyl-oxobutyrate |
| --- | --- | --- | --- | --- | --- |
| Concentration (mg/ml) | 3 | 2.5 | 0.04 | 0.007 | 3 |

*4.2. Vitamin A*

According to Figure 6 and Table 3, the pattern of changes in viabilities or IVCC of ‘Vit A in Ethanol (0.05 mg/ml)’ group is similar to the patterns of ‘Control’ and ‘Control+Ethanol’, while in other groups, the viabilities or IVCCs were very much lower than the corresponding control groups. In other words, any toxic effect of vitamin A is not evident in such concentration (0.05 mg/ml) and this concentration is chosen to be used in feed solutions.

**Table 5**. The final concentrations of the 15 metabolites that were used as (+1) level in Plackett-Burman design of experiment.

| **Name of metabolite** | **Concentration (mg/ml)** |
| --- | --- |
| Gln | 1.169 |
| Asn | 2.450 |
| Lys | 4.500 |
| Trp | 0.825 |
| Thr | 1.83 |
| Val | 2.840 |
| His | 1.300 |
| Vitamin B1 | 0.010 |
| Vitamin B6 | 0.010 |
| Thymidine | 3.000 |
| Deoxycytidine | 2.500 |
| Arachidonate | 0.040 |
| Deoxyguanosine | 0.007 |
| 3-methyl-oxobutyrate | 3.000 |
| Vitamin A | 0.050 |
