## Supplementary file S3 for "A metabolic network-based approach for developing feeding strategies for CHO cells to increase monoclonal antibody production"

* Corresponding authors

**S3 file:**

**Detailed statistical analysis of the results of Plackett-Burman designed experiments.**

**1. Introduction**

As mentioned in the main text of the manuscript, in order to validate the positive effect of the 15 metabolites on monoclonal antibody (mAb) production in CHO cells, a 20 run Plackett-Burman designed experiments were performed. Detailed changes in the viability of the cells and integral viable cell counts (IVCC) in every 20 groups will be shown here. Then, data regarding the fitting of the Hill function to the IVCC changes will be represented. In the end, the detailed statistical analysis of the results, which was performed using the Design-Expert software version 7 (Stat-EaseInc. Minneapolis, Minnesota, USA), will be discussed here.

**2. Results:**

**2.1. Changes in cell viability and IVCC**

Based on the PB design (as shown in Figure 1 of the main text of the manuscript), the effects of cell culture feeds on 20 groups were examined. Changes in the viability of the cells and IVCC in every 20 groups are shown in Figure S1.

A

Viability (%)

| Days | Group 1 | Group 2 | Group 3 | Group 4 | Group 5 | Group 6 | Group 7 | Group 8 | Group 9 | Group 10 | Group 11 | Group 12 | Group 13 | Group 14 | Group 15 | Group 16 | Group 17 | Group 18 | Group 19 | Group 20 |
| --- | --- | --- | --- | --- | --- | --- | --- | --- | --- | --- | --- | --- | --- | --- | --- | --- | --- | --- | --- | --- |
| 0 | 87.87 | 87.87 | 87.87 | 87.87 | 87.87 | 87.87 | 87.87 | 87.87 | 87.87 | 87.87 | 87.87 | 87.87 | 87.87 | 87.87 | 87.87 | 87.87 | 87.87 | 87.87 | 1 | 87.87 |
| 1 | 80 | 87.65 | 84.93 | 84.48 | 85.94 | 83.78 | 85.91 | 89.33 | 83.64 | 89.7 | 87.5 | 87.72 | 84.93 | 93.33 | 87.93 | 83.08 | 79.66 | 85.91 |  | 88.23 |
| 2 | 87.6 | 87.23 | 87.6 | 86.67 | 79.53 | 82.14 | 92.31 | 83.33 | 76.34 | 84.52 | 91.3 | 92.52 | 90.09 | 92.17 | 86.67 | 88 | 90.08 | 82.44 |  | 94.19 |
| 3 | 92.15 | 94.58 | 88.09 | 84.21 | 89.68 | 86.21 | 88.36 | 84.03 | 70.93 | 82.11 | 88.49 | 90.69 | 91.18 | 90.07 | 86.36 | 89.01 | 90.24 | 95.62 |  | 87.12 |
| 4 | 87.02 | 93.41 | 91.73 | 80.36 | 91.53 | 84.21 | 81.2 | 79.52 | 52.44 | 82.44 | 85.82 | 98.96 | 89.67 | 92.64 | 85.82 | 85.94 | 82.59 | 92.91 |  | 87.76 |
| 5 | 86.6 | 85.51 | 91.33 | 77.51 | 91.61 | 87.03 | 62.38 | 75.86 | 31.82 | 69.23 | 80 | 88.95 | 96.52 | 87.08 | 78.43 | 85.48 | 87.84 | 93.9 |  | 88.64 |
| 6 | 79.7 | 84.11 | 85.9 | 64.18 | 89.65 | 77.5 | 41.18 | 59.32 |  | 58.73 | 74.29 | 89.65 | 90.95 | 90.83 | 74.42 | 84.76 | 81.93 | 93.13 |  | 84.44 |
| 7 | 78.61 | 75 | 83.43 | 50.77 | 89.03 | 65.62 |  | 31.15 |  | 13.12 | 76.83 | 86.12 | 86.85 | 79.76 | 43.21 | 69.56 | 79.5 | 86.26 |  | 75.45 |
| 8 | 68.57 | 74.37 | 82.35 | 34.61 | 74.44 | 60.19 |  |  |  |  | 55.84 | 86.63 | 87.64 | 75.92 |  | 70.21 | 72.96 | 89.71 |  | 78.33 |
| 9 | 71.16 | 60.84 | 77.66 |  | 68.95 | 27.72 |  |  |  |  | 30.43 | 82.72 | 76.37 | 82.73 |  | 44.76 | 66.22 | 81.88 |  | 62.5 |
| 10 | 64.47 | 51.56 | 70.93 |  | 64 |  |  |  |  |  |  | 77.99 | 69.93 | 76.96 |  |  | 49.59 | 68.3 |  | 50.38 |
| 11 | 56.37 | 48.24 | 68.21 |  | 59.9 |  |  |  |  |  |  | 70.47 | 57.09 | 72.19 |  |  |  | 69.39 |  | 48.62 |
| 12 | 46.25 |  | 61.11 |  | 65.52 |  |  |  |  |  |  | 55.89 | 49.15 | 65.88 |  |  |  | 56.07 |  |  |
| 13 |  |  | 56.34 |  | 49.4 |  |  |  |  |  |  | 60.24 |  | 63.49 |  |  |  | 51.14 |  |  |
| 14 |  |  | 47.75 |  |  |  |  |  |  |  |  | 53.21 |  | 54.59 |  |  |  | 39.21 |  |  |
| 15 |  |  |  |  |  |  |  |  |  |  |  | 43.68 |  | 50 |  |  |  |  |  |  |

B

IVCC (million viable cells)

| Days | Group 1 | Group 2 | Group 3 | Group 4 | Group 5 | Group 6 | Group 7 | Group 8 | Group 9 | Group 10 | Group 11 | Group 12 | Group 13 | Group 14 | Group 15 | Group 16 | Group 17 | Group 18 | Group 19 | Group 20 |
| --- | --- | --- | --- | --- | --- | --- | --- | --- | --- | --- | --- | --- | --- | --- | --- | --- | --- | --- | --- | --- |
| 0 | 2.5 | 2.5 | 2.5 | 2.5 | 2.5 | 2.5 | 2.5 | 2.5 | 2.5 | 2.5 | 2.5 | 2.5 | 2.5 | 2.5 | 2.5 | 2.5 | 2.5 | 2.5 | 2.5 | 2.5 |
| 1 | 5.832 | 5.979 | 5.538 | 4.901 | 5.195 | 5.538 | 5.489 | 5.783 | 4.754 | 5.489 | 5.244 | 4.95 | 5.538 | 5.93 | 4.999 | 5.146 | 4.803 | 5.489 | 5.391 | 6.175 |
| 2 | 10.92 | 11.883 | 10.626 | 8.645 | 10.043 | 8.85 | 8.945 | 8.423 | 8.162 | 8.897 | 9.276 | 9.702 | 10.338 | 11.018 | 8.743 | 8.314 | 10.467 | 10.673 | 7.551 | 10.063 |
| 3 | 20.072 | 21.867 | 18.322 | 14.469 | 17.271 | 15.35 | 15.653 | 13.623 | 11.334 | 14.149 | 15.672 | 19.322 | 18.398 | 18.09 | 14.671 | 12.526 | 18.163 | 18.629 | 10.203 | 16.043 |
| 4 | 29.303 | 30.537 | 30.205 | 19.059 | 28.848 | 22.694 | 21.161 | 16.989 | 13.527 | 19.657 | 21.843 | 30.287 | 32.117 | 29.004 | 20.842 | 18.136 | 26.629 | 31.991 | 11.733 | 26.651 |
| 5 | 39.258 | 43.517 | 47.585 | 26.264 | 43.258 | 31.549 | 24.901 | 23.039 |  | 26.092 | 31.303 | 48.437 | 50.432 | 46.054 | 29.642 | 29.796 | 38.949 | 51.626 |  | 39.521 |
| 6 | 50.706 | 53.237 | 62.057 | 30.908 | 60.106 | 38.245 |  | 26.819 |  | 30.088 | 36.919 | 65.285 | 72.14 | 68.518 | 36.554 | 39.408 | 46.293 | 77.978 |  | 51.833 |
| 7 | 64.38 | 65.639 | 76.473 | 34.406 | 74.734 | 44.923 |  |  |  | 30.83 | 43.597 | 90.937 | 91.75 | 82.722 | 40.264 | 46.192 | 59.861 | 102.676 |  | 60.631 |
| 8 | 75.324 | 79.205 | 90.837 |  | 90.01 | 51.991 |  |  |  |  | 48.499 | 109.405 | 127.318 | 103.926 |  | 53.716 | 73.085 | 134.482 |  | 71.347 |
| 9 | 88.316 | 88.949 | 107.973 |  | 106.922 |  |  |  |  |  |  | 134.605 | 158.454 | 124.31 |  | 58.98 | 84.061 | 162.818 |  | 78.627 |
| 10 | 100.076 | 96.869 | 122.613 |  | 120.362 |  |  |  |  |  |  | 158.845 | 184.134 | 141.95 |  |  | 91.261 | 184.538 |  | 86.427 |
| 11 | 113.646 |  | 136.537 |  | 134.286 |  |  |  |  |  |  | 183.625 | 204.076 | 157.88 |  |  |  | 208.61 |  | 92.681 |
| 12 |  |  | 149.011 |  | 146.256 |  |  |  |  |  |  | 202.147 | 218.692 | 175.394 |  |  |  | 228.392 |  |  |
| 13 |  |  | 162.775 |  |  |  |  |  |  |  |  | 227.319 |  | 195.234 |  |  |  | 247.86 |  |  |
| 14 |  |  |  |  |  |  |  |  |  |  |  | 246.987 |  | 208.566 |  |  |  | 259.608 |  |  |
| 15 |  |  |  |  |  |  |  |  |  |  |  | 262.717 |  | 224.816 |  |  |  |  |  |  |

**Figure S1**. Changes in CHO cell viabilities (A) and integral viable cell count (IVCC) (B) of group 1-20. CHO cell viabilities have been shown by the percent of viable cells to the total number of cells. IVCC has been calculated by the cumulative addition of viable cell counts in millions for each day during CHO cell culture supplementation with 20 different feeds. The compositions of the feeds have been shown in Figure 1 in the manuscript.

**2.2. Hill function fitting**

In order to have a better representation of changes in the IVCC of each group, Hill function was fitted to changes in the IVCC of CHO cells during cell culture using the following equation:

$$\frac{\mathrm{IVCC}}{\mathrm{IVCC}_{max}}=\frac{D^{n}}{D^{n}+{D_{50}}^{n}}$$

IVCC_max_ is the maximum amount of IVCCs during cell culture, which equals the IVCC in the harvesting day. n represents the Hill coefficient. D represents the times that CHO cells are in cell culture medium (in days). D_50_ equals the day in which the cells reach a half amount of their maximum IVCC, which has been shown in Figure 2 in the main text of the manuscript. The groups that have bigger D_50_ values, have longer cell culture duration and therefore, better responses are expected in that group (e.g., group 12, 13, and 18). The details about the statistics of data fitting have been shown in Table S1.

**Table S1**. The statistics of fitting Hill function to the changes in the IVCC of each group in PB designed experiments. D_50_ equals the day in which the cells reach a half amount of their maximum IVCC, and n represents the Hill coefficient..

| **Groups** | **n** | **D_50_** | **R-square** |
| --- | --- | --- | --- |
| Group 1 | 2.43 | 1.95 | 0.9452 |
| Group 2 | 3.59 | 5.22 | 0.9782 |
| Group 3 | 3.59 | 5.94 | 0.983 |
| Group 4 | 3.40 | 4.20 | 0.9732 |
| Group 5 | 3.85 | 5.89 | 0.9843 |
| Group 6 | 3.66 | 4.71 | 0.9760 |
| Group 7 | 3.58 | 3.51 | 0.9684 |
| Group 8 | 3.15 | 3.77 | 0.9529 |
| Group 9 | 2.81 | 2.61 | 0.9579 |
| Group 10 | 3.60 | 3.85 | 0.9604 |
| Group 11 | 3.62 | 4.58 | 0.9770 |
| Group 12 | 4.05 | 6.50 | 0.9827 |
| Group 13 | 4.89 | 6.44 | 0.9815 |
| Group 14 | 3.73 | 6.25 | 0.9844 |
| Group 15 | 3.94 | 4.33 | 0.9633 |
| Group 16 | 4.22 | 5.15 | 0.9741 |
| Group 17 | 3.84 | 5.29 | 0.9707 |
| Group 18 | 4.36 | 6.38 | 0.9893 |
| Group 19 | 3.56 | 5.61 | 0.9733 |
| Group 20 | 3.73 | 5.26 | 0.9856 |

***2.3. Response 1: IVCC***

In order to have significant modeling of this response, the metabolites with less than 3% contribution to the response have not been modeled.

ANOVA for selected factorial model:

Analysis of variance table (Partial sum of squares - Type III):

| **Source** | **Sum of**  **Squares** | **Mean**  **Square** | **F-Value** | **p-value**  **Prob > F** |  |
| --- | --- | --- | --- | --- | --- |
| **Model** | 80362.97 | 13393.83 | 3.21 | 0.0367 | ***significant*** |
| Asparagine | 4733.07 | 4733.07 | 1.14 | 0.3059 |  |
| Lysine | 6323.39 | 6323.39 | 1.52 | 0.2398 |  |
| Tryptophan | 17836.51 | 17836.51 | 4.28 | 0.0590 |  |
| Threonine | 19335.43 | 19335.43 | 4.64 | 0.0506 |  |
| Retinol (Vitamin A) | 7203.10 | 7203.10 | 1.73 | 0.2113 |  |
| Arachidonate | 24931.47 | 24931.47 | 5.98 | 0.0294 |  |
| *Residual* | 54160.55 | 4166.20 |  |  |  |
| *Cor Total* | 1.345E+005 |  |  |  |  |

The Model F-value is 3.21, and therefore, the model is significant. In other words, there is only a 3.67% chance that the results could occur due to noise.

The "Prob > F" value of ‘P-T’ is significant (less than 0.0500).

| Std. Dev. | 64.55 |
| --- | --- |
| Mean | 103.14 |
| C.V. % | 62.58 |
| PRESS | 1.282E+005 |
| R-Squared | 0.5974 |
| *Adj R-Squared* | 0.4116 |
| *Pred R-Squared* | 0.0471 |
| ***Adeq Precision*** | 6.036 |

The "*Pred R-Squared*" of 0.0471 is not as close to the "*Adj R-Squared*" of 0.4116. This may indicate a large block effect in our model and/or data. Model reduction, response transformation, outliers, etc. might be helpful in this regard.

"***Adeq Precision***" measures the signal to noise ratio. A ratio greater than 4 is desirable. Here, a ratio of 6.036 indicates an adequate signal. This model can be used to navigate the design space.

Design-Expert calculates the model coefficients in the final equation using regression estimates. For orthogonal, balanced designs, the model coefficients will be equal to one half the effects, which come from the difference between the average response at the high level and the average response at the low level.

**Final Equation:**

**IVCC = +103.14 + 15.38 * Asparagine + 17.78 * Lysine - 29.86 * Tryptophan + 31.09 * Threonine + 18.98 * Vitamin A +35.31 * Arachidonate**

It has to be noted that, the ANOVA modeling with all 15 metabolites was not significant. Therefore, we performed a sensitivity analysis, by restricting the model to the metabolites which are contributed to the results more than 0.5, 1, 1.5, 2, 2.5, 3, 3.5, 4, 4.5, and 5%. The two metabolites that we chose for performing RSM (threonine and arachidonate) persistently in all cases have a positive and significant impact on the responses. Therefore, we reported the significant modeling results with a minimum amount of restriction, which was omitting the metabolites with contributions of less than 3%. The effect list and the contribution of the metabolites have been listed in Table S2.

**Table S2**. The effect list and the contribution of the metabolites to the first response (IVCC).

| **Metabolites** | **Effects** | **Sum of Squares** | **Contributions (%)** |
| --- | --- | --- | --- |
| Gln | -15.1653 | 1149.93 | 0.854818 |
| Asn | 30.7671 | 4733.07 | 3.5184 |
| Lys | 35.5623 | 6323.39 | 4.70058 |
| Trp | -59.7269 | 17836.5 | 13.259 |
| Thr | 62.1859 | 19335.4 | 14.3733 |
| Val | 23.3415 | 2724.13 | 2.02502 |
| His | -16.5915 | 1376.39 | 1.02316 |
| Vit B1 | 9.0593 | 410.355 | 0.305043 |
| Vit B6 | -12.5775 | 790.968 | 0.587977 |
| Thymidine | -26.1235 | 3412.19 | 2.5365 |
| Deoxy-cytidine | -2.8039 | 39.3093 | 0.0292211 |
| 3-methyl-oxobutyrate | -0.5189 | 1.34629 | 0.00100078 |
| Deoxy-guanosine | -14.3861 | 1034.8 | 0.769233 |
| Vit A | 37.9555 | 7203.1 | 5.35453 |
| Arachidonate | 70.6137 | 24931.5 | 18.5332 |

***2.4. Response 2: Total mAb expression***

In order to have significant modeling of this response, the metabolites with less than 3% contribution to the response have not been modeled.

ANOVA for selected factorial model:

Analysis of variance table (Partial sum of squares - Type III):

| **Source** | **Sum of**  **Squares** | **Mean**  **Square** | **F-Value** | **p-value**  **Prob > F** |  |
| --- | --- | --- | --- | --- | --- |
| **Model** | 1.307E+005 | 16334.28 | 3.27 | 0.0363 | ***significant*** |
| Glutamine | 3489.49 | 3489.49 | 0.70 | 0.4211 |  |
| Asparagine | 4315.57 | 4315.57 | 0.86 | 0.3726 |  |
| Lysine | 4107.85 | 4107.85 | 0.82 | 0.3839 |  |
| Tryptophan | 40204.75 | 40204.75 | 8.05 | 0.0162 |  |
| Threonine | 39259.39 | 39259.39 | 7.86 | 0.0172 |  |
| Valine | 3305.25 | 3305.25 | 0.66 | 0.4332 |  |
| Retinol (Vitamin A) | 3595.87 | 3595.87 | 0.72 | 0.4143 |  |
| Arachidonate | 32396.05 | 32396.05 | 6.49 | 0.0272 |  |
| *Residual* | 54950.27 | 4995.48 |  |  |  |
| *Cor Total* | 1.856E+005 |  |  |  |  |

The Model F-value is 3.27, and therefore, the model is significant. In other words, there is only a 3.63% chance that the results could occur due to noise.

The "Prob > F" value of ‘D-D’, ‘E-E’, and ‘P-T’ are significant (less than 0.0500).

| Std. Dev. | 70.68 |
| --- | --- |
| Mean | 278.70 |
| C.V. % | 25.36 |
| PRESS | 1.817E+005 |
| R-Squared | 0.7040 |
| *Adj R-Squared* | 0.4887 |
| *Pred R-Squared* | 0.0214 |
| ***Adeq Precision*** | 5.946 |

The "*Pred R-Squared*" of 0.0214 is not as close to the "*Adj R-Squared*" of 0.4887. This may indicate a large block effect in our model and/or data. Model reduction, response transformation, outliers, etc. might be helpful in this regard.

"***Adeq Precision***" measures the signal to noise ratio. A ratio greater than 4 is desirable. Here, a ratio of 5.946 indicates an adequate signal. This model can be used to navigate the design space.

Design-Expert calculates the model coefficients in the final equation using regression estimates. For orthogonal, balanced designs, the model coefficients will be equal to one half the effects, which come from the difference between the average response at the high level and the average response at the low level.

**Final Equation: Total mAb expression = +278.70 - 13.21 * Glutamine + 14.69 * Asparagine + 14.33 * Lysine - 44.84 * Tryptophan + 44.31 * Threonine + 12.86 * Valine + 13.41 * VitaminA + 40.25 * Arachidonate**

The same as the first response, we performed a sensitivity analysis by restricting the model to the metabolites which are contributed to the results more than 0.5, 1, 1.5, 2, 2.5, 3, 3.5, 4, 4.5, and 5%. The two metabolites that we chose for performing RSM (threonine and arachidonate) persistently in all cases have a positive and significant impact on the responses. Therefore, we reported the significant modeling results with a minimum amount of restriction, which was omitting the metabolites with contributions of less than 3%. The effect list and the contribution of the metabolites have been listed in Table S3.

**Table S3**. The effect list and the contribution of the metabolites to the second response (mAb expression).

| **Metabolites** | **Effects** | **Sum of Squares** | **Contributions (%)** |
| --- | --- | --- | --- |
| Gln | -26.4177 | 3489.49 | 1.87986 |
| Asn | 29.3788 | 4315.57 | 2.32489 |
| Lys | 28.663 | 4107.85 | 2.21299 |
| Trp | -89.6713 | 40204.7 | 21.6592 |
| Thr | 88.6108 | 39259.4 | 21.1499 |
| Val | 25.7109 | 3305.25 | 1.78061 |
| His | 3.02813 | 45.8478 | 0.0246992 |
| Vit B1 | 3.16493 | 50.0838 | 0.0269813 |
| Vit B6 | -21.8779 | 2393.21 | 1.28927 |
| Thymidine | -8.44715 | 356.772 | 0.192201 |
| Deoxy-cytidine | 1.27241 | 8.09511 | 0.00436101 |
| 3-methyl-oxobutyrate | 9.81466 | 481.637 | 0.259469 |
| Deoxy-guanosine | -7.94308 | 315.463 | 0.169947 |
| Vit A | 26.8174 | 3595.87 | 1.93717 |
| Arachidonate | 80.4935 | 32396 | 17.4525 |

**3. Discussion**

According to the results of both responses, tryptophan, threonine, and arachidonate have a p-value equal to, or less than, 0.05, which means that these metabolites have significant effects on CHO cells. Based on both final equations, threonine and arachidonate have positive effects and tryptophan has a negative effect on the responses. Because we want to improve mAb production in CHO cells, we did not choose tryptophan for the next step (designing with RSM). In total, some of the metabolites have shown negative effects on mAb production or IVCC. This can be caused by the toxic effects of the metabolite in the level of concentration that has been used for CHO cells.
