## Supplementary file S4 for "A metabolic network-based approach for developing feeding strategies for CHO cells to increase monoclonal antibody production"

* Corresponding authors

**S4 file:**

**Detailed statistical analysis of the results of central composite designed experiments.**

**1. Introduction**

As mentioned in the main text of the manuscript, based on the results of Plackett-Burman designed experiments, the “positive” effects of the addition of arachidonate and threonine to cell culture feeds for increasing the monoclonal antibody (mAb) production was revealed. Then, response surface mythology (RSM) was used to find the best levels of concentrations of these two metabolites, by performing central composite design (CCD) experiments. CCD is the most commonly used method in RSM (1). It builds a quadratic model for the response, while using a three-level factorial experiment. CCD consists of a 2^k^ cube points (coded by +1 and -1), augmented by 2*k star points (coded by +α and -α), and a number of centre points (coded by 0). CCD can be rotatable by choosing α (2). Here, we have two factors (k=2), and three center points (Figure S1). As shown in figure S1, the α equals √2=1.41.

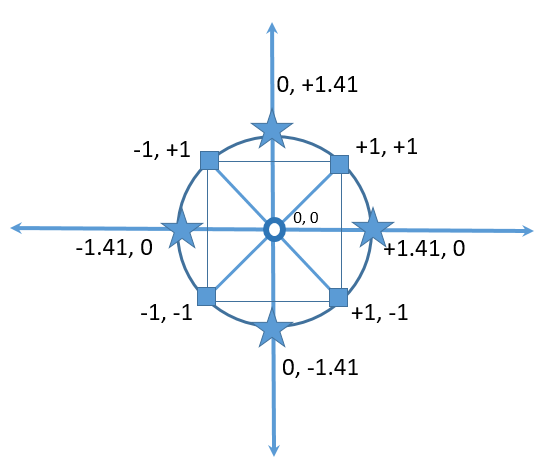

**Figure S1**. The cube (shown by ◼), star (shown by ★), and center (shown by 🞈) points of a CCD for two factors. ”

A detailed statistical analysis of the results, which was performed using the Design-Expert software version 7 (Stat-EaseInc. Minneapolis, Minnesota, USA), has been discussed here.

**2. Results:**

**2.1. Changes in cell viability and IVCC**

Based on the CCD design (as shown in Figure 3 of the main text of the manuscript), the effects of cell culture feeds on 11 groups were examined. Changes in the viability of the cells and IVCC in every 11 groups are shown in Figure S2.

A

Viability (%)

| Days | F01 | F02 | F03 | F04 | F05 | F06 | F07 | F08 | F09 | F10 | F11 |
| --- | --- | --- | --- | --- | --- | --- | --- | --- | --- | --- | --- |
| 0 | 82 | 82 | 82 | 82 | 82 | 82 | 82 | 82 | 82 | 82 | 82 |
| 1 | 80 | 75.75 | 71.19 | 75.41 | 79.41 | 78.69 | 73.97 | 74.65 | 84.13 | 83.33 | 83.72 |
| 2 | 85.71 | 80.13 | 79.39 | 83.56 | 83.05 | 81.76 | 83.78 | 86.15 | 81.48 | 85.43 | 90.48 |
| 3 | 79.54 | 78.45 | 76.52 | 82.25 | 82.35 | 82.96 | 84.91 | 84.98 | 78.77 | 80.33 | 87.36 |
| 4 | 78.33 | 62.12 | 63.79 | 76.29 | 73.68 | 82.22 | 85.48 | 79.02 | 75 | 75.86 | 77.78 |
| 5 | 67.11 | 51.25 | 56.38 | 67.06 | 67.1 | 75.25 | 78.68 | 70.68 | 52.67 | 67.14 | 73.91 |
| 6 | 59.375 | 44.03 | 43.78 | 61.22 | 55.67 | 70.35 | 69.2 | 63.83 | 41.22 | 63.13 | 63.24 |
| 7 | 51.98 |  |  | 50.32 | 40.72 | 60.71 | 64.7 | 51.1 |  | 62.63 | 61.39 |
| 8 | 50 |  |  | 43.36 |  | 56.3 | 61.05 | 50.34 |  | 57.86 | 62.05 |
| 9 | 43.6 |  |  |  |  | 45.45 | 53.62 | 35.81 |  | 44.62 | 59.22 |
| 10 |  |  |  |  |  |  | 55.08 |  |  |  | 52.74 |
| 11 |  |  |  |  |  |  | 42.19 |  |  |  | 48.04 |

B

IVCC (million viable cells)

| Days | F01 | F02 | F03 | F04 | F05 | F06 | F07 | F08 | F09 | F10 | F11 |
| --- | --- | --- | --- | --- | --- | --- | --- | --- | --- | --- | --- |
| 0 | 2.5 | 2.5 | 2.5 | 2.5 | 2.5 | 2.5 | 2.5 | 2.5 | 2.5 | 2.5 | 2.5 |
| 1 | 5.244 | 4.95 | 4.558 | 4.754 | 6.469 | 4.852 | 5.146 | 5.097 | 5.097 | 4.705 | 6.028 |
| 2 | 7.692 | 7.854 | 7.054 | 7.682 | 8.821 | 7.756 | 8.122 | 7.785 | 8.265 | 8.785 | 8.764 |
| 3 | 12.627 | 12.131 | 11.19 | 12.0633 | 14.085 | 12.4222 | 12.7515 | 12.8375 | 12.5288 | 13.4042 | 14.451 |
| 4 | 17.4279 | 14.222 | 14.964 | 15.8373 | 18.369 | 16.1962 | 18.1575 | 17.8227 | 15.8948 | 17.8922 | 18.735 |
| 5 | 22.4279 | 16.272 | 17.614 | 18.6873 | 20.919 | 19.9962 | 23.5075 | 22.5227 | 18.6948 | 20.2422 | 22.985 |
| 6 | 25.5059 | 17.865 | 19.801 | 21.9273 | 23.3693 | 23.2632 | 28.9075 | 25.6075 | 20.1528 | 23.9412 | 26.144 |
| 7 | 28.9774 |  |  | 24.0208 | 25.1713 | 25.9662 | 32.697 | 28.072 |  | 27.2272 | 29.642 |
| 8 | 31.4284 |  |  | 25.4173 |  | 27.8757 | 36.003 | 30.181 |  | 29.8492 | 32.5775 |
| 9 | 34.0044 |  |  |  |  | 29.5557 | 39.111 | 31.665 |  | 31.9492 | 35.5455 |
| 10 |  |  |  |  |  |  | 43.011 |  |  |  | 38.7255 |
| 11 |  |  |  |  |  |  | 45.4005 |  |  |  | 41.2625 |

**Figure S2**. Changes in CHO cell viabilities (A) and integral viable cell count (IVCC) (B) of group 1-11, supplementation with 11 different feeds (F01-F11).. CHO cell viabilities have been shown by the percent of viable cells to the total number of cells. IVCC has been calculated by the cumulative addition of viable cell counts in millions for each day during CHO cell culture supplementation with 11 different feeds. The compositions of the feeds have been shown in Figure 3 in the manuscript.

**2.2. Hill function fitting**

In order to have a better representation of changes in the IVCC of each group, Hill function was fitted to changes in the IVCC of CHO cells during cell culture using the following equation:

$$\frac{\mathrm{IVCC}}{\mathrm{IVCC}_{max}}=\frac{D^{n}}{D^{n}+{D_{50}}^{n}}$$

IVCC_max_ is the maximum amount of IVCCs during cell culture, which equals the IVCC in the harvesting day. n represents the Hill coefficient. D represents the times that CHO cells are in cell culture medium (in days). D_50_ equals the day in which the cells reach a half amount of their maximum IVCC, which has been shown in Figure 2 in the main text of the manuscript. The groups that have bigger D_50_ values, have longer cell culture duration and therefore, better responses are expected in that group (e.g., group 12, 13, and 18). The details about the statistics of data fitting have been shown in Table S1.

**Table S1**. The statistics of fitting Hill function to the changes in the IVCC of each group in CCD experiments. D_50_ equals the day in which the cells reach a half amount of their maximum IVCC, and n represents the Hill coefficient.

| **Groups** | **n** | **D_50_** | **R-square** |
| --- | --- | --- | --- |
| Group 1 | 3.02 | 4.656 | 0.9757 |
| Group 2 | 2.78 | 2.986 | 0.9645 |
| Group 3 | 3.09 | 3.415 | 0.9530 |
| Group 4 | 2.92 | 3.966 | 0.9708 |
| Group 5 | 2.82 | 3.443 | 0.9667 |
| Group 6 | 2.89 | 4.375 | 0.9753 |
| Group 7 | 2.97 | 5.579 | 0.9795 |
| Group 8 | 3.11 | 4.374 | 0.9794 |
| Group 9 | 2.99 | 3.165 | 0.9615 |
| Group 10 | 2.74 | 4.431 | 0.9741 |
| Group 11 | 2.56 | 5.208 | 0.9748 |

***2.3. Response 1: IVCC***

ANOVA for Response Surface 2FI Model

A = Threonine

B = Arachidonate

| **Source** | **Sum of**  **Squares** | **Mean**  **Square** | **F-Value** | **p-value**  **Prob > F** |  |
| --- | --- | --- | --- | --- | --- |
| **Model** | 530.02 | 176.67 | 5.07 | 0.0355 | ***significant*** |
| **A-A** | 0.051 | 0.051 | 1.472E-003 | 0.9705 |  |
| **B-B** | 520.69 | 520.69 | 14.94 | 0.0062 |  |
| **AB** | 9.27 | 9.27 | 0.27 | 0.6218 |  |
| *Residual* | 243.90 | 34.84 |  |  |  |
| *Lack of Fit* | 202.01 | 40.40 | 1.93 | 0.3757 | *not significant* |
| *Pure Error* | 41.89 | 20.94 |  |  |  |
| *Cor Total* | 773.92 |  |  |  |  |

The Model F-value is 5.07, and therefore, the model is significant. In other words, there is only a 3.55% chance that the results could occur due to noise.

The "Prob > F" value of ‘B’ or arachidonate is significant (less than 0.0500).

The "*Lack of Fit F-value*" is 1.93, *i.e.*, it is not significant relative to the pure error. There is a 37.57% chance that a "*Lack of Fit F-value*" could occur due to noise. Non-significant lack of fit is good -- we want the model to fit.

| Std. Dev. | 5.90 |
| --- | --- |
| Mean | 29.29 |
| C.V. % | 20.15 |
| PRESS | 820.09 |
| R-Squared | 0.6848 |
| Adj R-Squared | 0.5498 |
| ***Pred R-Squared*** | -0.0597 |
| ***Adeq Precision*** | 6.411 |

A negative "***Pred R-Squared***" implies that the overall mean is a better predictor of the response than the current model.

"***Adeq Precision***" measures the signal to noise ratio. A ratio greater than 4 is desirable. Here, a ratio of 6.411 indicates an adequate signal. This model can be used to navigate the design space.

**Final Equation: IVCC = +29.29 + 0.080 * A + 8.07 * B + 1.52 * AB**

***2.4. Response 2: Total mAb expression***

ANOVA for Response Surface 2FI Model

A = Threonine

B = Arachidonate

| **Source** | **Sum of**  **Squares** | **Mean**  **Square** | **F-Value** | **p-value**  **Prob > F** |  |
| --- | --- | --- | --- | --- | --- |
| **Model** | 4427.96 | 1475.99 | 9.02 | 0.0084 | ***significant*** |
| **A-A** | 10.04 | 10.04 | 0.061 | 0.8114 |  |
| **B-B** | 4105.35 | 4105.35 | 25.10 | 0.0015 |  |
| **AB** | 312.57 | 312.57 | 1.91 | 0.2094 |  |
| *Residual* | 1145.12 | 163.59 |  |  |  |
| *Lack of Fit* | 794.59 | 158.92 | 0.91 | 0.5989 | *not significant* |
| *Pure Error* | 350.53 | 175.27 |  |  |  |
| *Cor Total* | 5573.08 |  |  |  |  |

The Model F-value is 9.02, and therefore, the model is significant. In other words, there is only a 0.84% chance that the results could occur due to noise.

The "Prob > F" value of ‘B’ or arachidonate is significant (less than 0.0500).

The "Lack of Fit F-value" is 0.91, *i.e.*, it is not significant relative to the pure error. There is a 59.89% chance that a "*Lack of Fit F-value*" could occur due to noise. Non-significant lack of fit is good -- we want the model to fit.

| Std. Dev. | 12.79 |
| --- | --- |
| Mean | 168.14 |
| C.V. % | 7.61 |
| PRESS | 3413.82 |
| R-Squared | 0.7945 |
| ***Adj R-Squared*** | 0.7065 |
| ***Pred R-Squared*** | 0.3874 |
| ***Adeq Precision*** | 8.382 |

The "***Pred R-Squared***" of 0.3874 is not as close to the "***Adj R-Squared***" of 0.7065 as one might normally expect. This may indicate a large block effect or a possible problem with our data. Things to consider are model reduction, response transformation, outliers, etc.

"***Adeq Precision***" measures the signal to noise ratio. A ratio greater than 4 is desirable. Here, a ratio of 8.382 indicates an adequate signal. This model can be used to navigate the design space.

**Final Equation: Total mAb expression = +168.14 + 1.12 * A + 22.65 * B + 8.84 * AB**

***2.5. Predicted optimum points***

Supposing the range of concentrations of both threonine and arachidonate to be changing between ‘-1’ and ‘+1’ levels, a set of optimum points are also predicted by the software, as follows:

| Number | Threonine level | Arachidonate level | IVCC | Total mab expression |
| --- | --- | --- | --- | --- |
| 1 | +1.00 | +1.00 | 38.9653 | 200.749 |
| 2 | +0.12 | +1.00 | 37.5538 | 191.977 |

**3. Discussion**

According to the results, arachidonate and threonine are positively interacting with each other to increase both IVCC and mAb expression, as it has been shown that the first predicted optimum point is in ’+1’ level. However, the impact of threonine on IVCC and mAb expression is not very large (it was not significant in modeling). This issue can be seen on the predicted optimum points because decreasing the level of threonine concentration form ‘+1’ to ‘+0.12’ has only a minor decreasing effect on IVCC and mAb expression.
